# KDR Signaling in Muscle Stem Cells Promotes Asymmetric Division and Progenitor Generation for Efficient Regeneration

**DOI:** 10.1101/2022.06.27.497734

**Authors:** William Chen, Yu Xin Wang, Morten Ritso, Theodore J. Perkins, Michael A. Rudnicki

## Abstract

The regulation of muscle stem cell (MuSC) asymmetric division plays an essential role in controlling the growth and repair of skeletal muscle. We discover kinase domain receptor (KDR) as a positive modulator of MuSC asymmetric division using an *in-niche* high-content screen and confirmed its expression in satellite cells by ddPCR and immunofluorescence. Knockdown of KDR significantly reduces the numbers of asymmetric divisions, whereas ligand stimulation of KDR increases the numbers of asymmetric divisions. KDR signaling is impaired in dystrophin-deficient satellite cells and requires a polarized cell environment established by the dystrophin glycoprotein complex (DGC) to direct asymmetric division. Mice lacking KDR in MuSCs exhibit reduced numbers of satellite cells due to precocious differentiation, and deficits in regeneration consistent with impaired asymmetric division and reduced generation of progenitors. Therefore, our experiments identify KDR signaling as playing an essential role in MuSC function in muscle regeneration.

**HIGHLIGHTS:**

- KDR and VEGFA are expressed in satellite cells
- Ligand activated KDR stimulates asymmetric satellite stem cell division
- KDR signaling requires the presence of the DGC
- KDR-deficient satellite cells give rise to reduced numbers of progenitors

**eTOC blurb:** Chen et al., performed a chemical screen using a novel screening platform to identify modulators of muscle stem cell asymmetric division. They discovered that KDR signalling requires the presence of the dystrophin associated glycoprotein complex and is an important regulator of muscle stem cell asymmetric division.

## INTRODUCTION

Muscle satellite cells are required for the growth and regeneration of skeletal muscle (Yin et al., 2013). These cells are identified by the expression of transcription factor *Pax7* and express a number of membrane markers including syndecan-4 (Sdc4), vascular Cell adhesion molecule 1 (VCAM1), alpha-5 integrin (Itga7), notch receptors, calcintonin receptor (CalcR), frizzled 7 (Fzd7), and epiderman growth factor receptor (EGFR) (Baghdadi et al., 2018a; Cornelison et al., 2004; Fukada et al., 2007; Gnocchi et al., 2009; Le Grand et al., 2009; Kann and Krauss, 2019; Seale et al., 2000; Wang et al., 2019; Yin et al., 2013; Zhang et al., 2019). Satellite cells are a heterogeneous population where the majority are a short-term repopulating cells (Kuang et al., 2007), with a minority subset of long-term self-renewing stem cells that give rise to committed progenitors through asymmetric cell divisions (Gurevich et al., 2016; Kuang et al., 2007; Rocheteau et al., 2012). Notably, the long-term self-renewing stem cells do not express the myogenic transcription factor Myf5, which distinguishes stem cells from short-term self-renewing cells and committed progenitors (Kuang et al., 2007).

The regulatory pathways that regulate the choice between MuSC symmetric versus asymmetric division have a significant impact on the efficiency of muscle regeneration. Wnt7a/Fzd7 stimulation of the planar cell polarity pathway drives MuSC symmetric expansion increasing satellite cell numbers and enhancing regeneration (Le Grand et al., 2009). By contrast, increased JAK2/STAT3 signalling during aging results in reduced numbers of MuSC by inhibiting symmetric divisions (Price et al., 2014; Tierney et al., 2014). Similarly, activation of p38α MAPK signaling in MuSC promotes asymmetric division, commitment to differentiation in ageing, and loss of regenerative potential (Bernet et al., 2014; Cosgrove et al., 2014). Recently, we identified EGFR/Aurka signaling as regulating asymmetric MuSC divisions and that activation of this pathway significantly ameliorated disease progression in *mdx* mice, a mouse model for Duchenne’s Muscular Dystrophy (DMD), by rescuing the loss of asymmetric divisions in dystrophin deficient satellite cells (Wang et al., 2019). Therefore, the control of MuSC asymmetric division plays a key role in regulating the efficiency of muscle regeneration in health and in disease.

Polarization of the intrinsic and extrinsic cell environment is essential for maintaining satellite stem cell identity and quiescence (Baghdadi et al., 2018b, 2018a; Bernet et al., 2014; Chang et al., 2018; Chen et al., 2020; Cosgrove et al., 2014; Crist et al., 2012; Dumont et al., 2015; Le Grand et al., 2009; Haller et al., 2017; Kuang et al., 2007; Morrée et al., 2017; Price et al., 2014; Shea et al., 2010; Verma et al., 2018; Zhang et al., 2019). The immediate satellite cell niche is a polarized environment with the satellite cell situated between the basal lamina and myofiber sarcolemma (Yin et al., 2013). This polarization aids the satellite cell in providing cues to divide asymmetrically in an apical-basal fashion (Kuang et al., 2007; Wang et al., 2019).

The intrinsic polarized satellite cell environment is established in part by the dystrophin-associated glycoprotein complex (DGC) (Dumont et al., 2015). Together, the extrinsic and intrinsic polarized cues and environment are necessary for the asymmetric segregation of transcription factors and proteins required for asymmetric division to occur (Chang et al., 2018; Conboy and Rando, 2002; Dumont and Rudnicki, 2017; Price et al., 2014; Shea et al., 2010; Shinin et al., 2006; Wang et al., 2019). The loss of cell polarity, such as in DMD, results in reduced numbers of asymmetric divisions, reduced generation of progenitors, and a decline in muscle regenerative capacity (Dumont et al., 2015).

Multiple reports have provided evidence that an endothelial-like gene expression signature is present in satellite cells (Angelis et al., 1999; Cosgrove et al., 2009; Giordani et al., 2019; Goel et al., 2017; Mayeuf-Louchart et al., 2014; Motoike et al., 2003). Interestingly, Pax7-expressing satellite cells are derived from the same mesodermal multipotent Pax3^+^ population that gives rise to CD31^+^ endothelial cells (Mayeuf-Louchart et al., 2014; Motoike et al., 2003). Kinase Domain Receptor (KDR), also known as VEGFR2 and Flk1, is a receptor tyrosine kinase activated by binding of VEGFA, VEGFC and VEGFD, that plays an essential role in angiogenesis, vascular development, vascular permeability, and embryonic hematopoiesis (Apte et al., 2019). Notably, VEGFA ligand is abundantly expressed within the muscle by nearly all components of the satellite cell niche including myofibers, satellite cells, endothelial cells, and fibroadipogenic cells (Blackburn et al., 2019; Christov et al., 2007; Micheli et al., 2020; Rissanen et al., 2002; Verma et al., 2018; Wagatsuma et al., 2006).

Here we employed an *ex vivo* high content MuSC screen and identified KDR signaling as a modulator of asymmetric division. siRNA mediated knockdown of KDR significantly inhibits asymmetric divisions. Conversely, ligand stimulation of KDR increases the numbers of muscle stem cell asymmetric divisions. This pathway is impaired in *mdx* mice and cannot be rescued by ligand treatment. Impairment of KDR-induced muscle stem cell asymmetric divisions result in a cascading effect on muscle regeneration, leading to fewer progenitors in response to injury. Therefore, we conclude the KDR pathway is a critical component of stem cell function, and this process is necessary for both muscle regeneration and maintenance.

## RESULTS

### Identification of KDR as a modulator of muscle stem cell asymmetric division

The satellite cell is extremely sensitive to its microenvironment. Changes to the satellite cell niche lead to changes in satellite cell identity in merely hours (Machado et al., 2017; Velthoven et al., 2017). Therefore, we devised a novel image-based high content analysis platform to quantify satellite stem cell fate decisions using *ex vivo* culture of single myofibers to preserve the satellite cell niche. We combined the *Myf5-Cre* (Tallquist et al., 2000) and *R26R-eYFP* (Srinivas et al., 2001) alleles to visualize stem cell fate decisions as previously described (Kuang et al., 2007; Wang et al., 2019) (Figure 1A). This imaging approach allows for lineage tracing of muscle stem cells (Myf5^-^ and eYFP^-^) that commit (Myf5^+^ and eYFP^+^) to the myogenic lineage.

**Figure 1.**
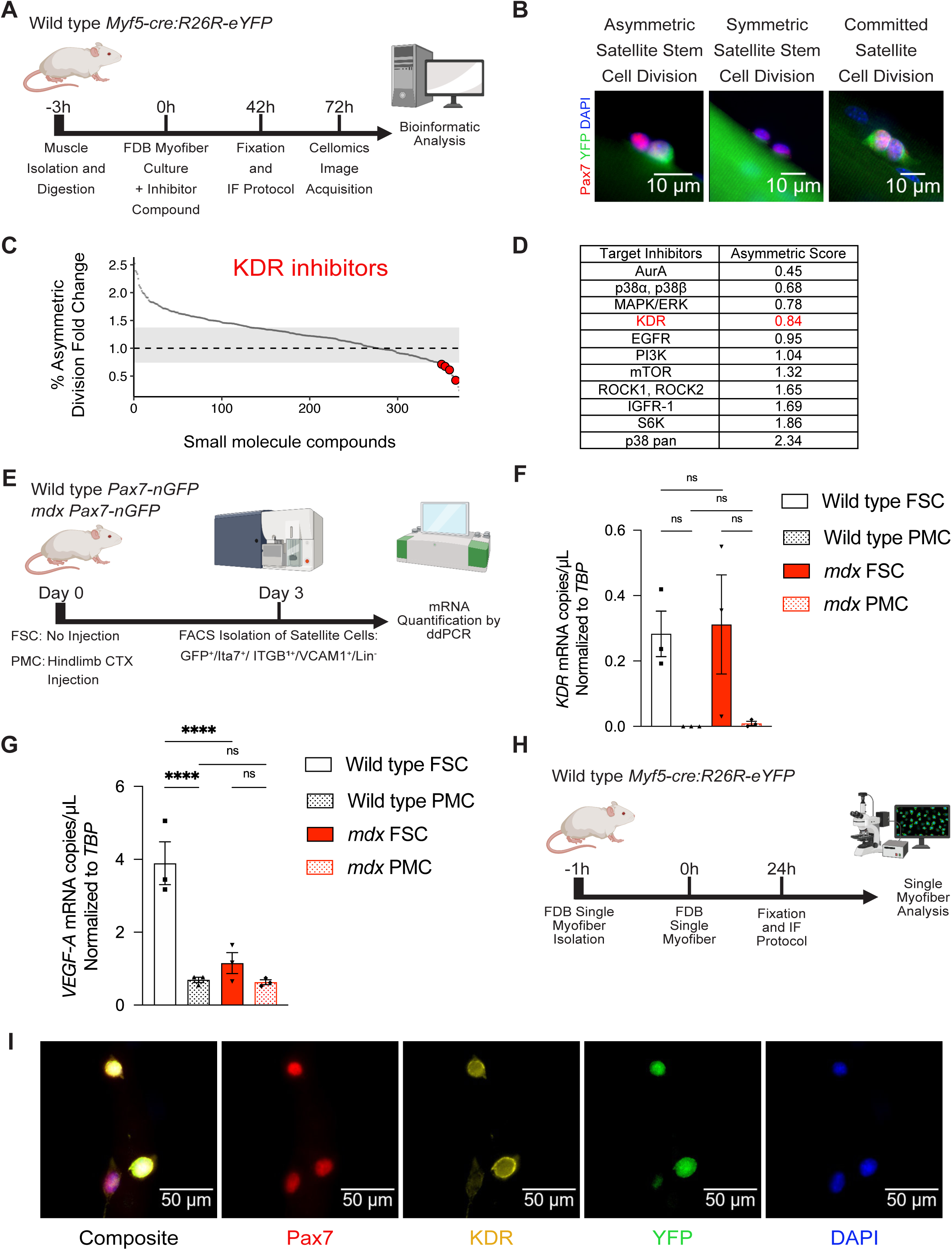
KDR is expressed in satellite cells. (A) Schematic of high content analysis workflow. Single myofibers are cultured in growth medium (vehicle) supplemented with inhibitor compounds. EGF and Wnt7a ligand treatment are used as positive controls for asymmetric and symmetric divisions respectively. Fixation is performed at 42h for quantification of satellite cell divisions. Image acquisition is performed by the Cellomics platform. Findings are analyzed using a custom bioinformatic MATLAB-based script to quantify satellite cell divisions. See also Figure S1A-B. (B) Representative immunofluorescence (IF) images of satellite cell divisions derived from wild type *Myf5-Cre:R26R-eYF*P mice. Satellite cells are identified by Pax7^+^ expression (red). Stem cells are eYFP^-^, whereas activated satellite cells are eYFP^+^ (green). Nuclei were stained with DAPI (blue). (C) High content analysis of satellite stem cell divisions. Percent fold change in asymmetric division compared to vehicle treatment is quantified. Grey area limits for asymmetric fold change is determined by positive controls EGF and Wnt7a. Each grey dot is a single inhibitor compound. Top KDR inhibitor compounds are shown in red. (D) Asymmetric scoring of inhibitor targets based on frequency of hits and IC50 of compounds screened compared to concentration screened (1μM). Values lower than 1 report negative effectors of asymmetric division, whereas values higher than 1 report positive effectors of asymmetric division. (E) Schematic of satellite cell isolation by FACs experimental design. Positive markers for satellite cells include GFP, α7-integrin (ITGA), β1-integrin (ITGB1), and VCAM1. Negative lineage markers (Lin^-^) include CD45, CD11b, Sca-1 and CD31. See also Figure S1C. (F) *KDR* expression by ddPCR from FACS isolated Freshly sorted Satellite Cells (FSC) and Proliferating Myogenic Precursor Cells (PMC) from wild type and *mdx* hindlimb. Data is presented as mean. Error bars are ± SEM (n=3). See also Figure S1D. (G) *VEGFA* expression by ddPCR from FACs isolated Freshly sorted Satellite Cells (FSC) and Proliferating Myogenic Precursor Cells (PMC) from wild type and mdx hindlimb. Data is presented as mean. Error bars are ± SEM (n=3). See also Figure S1E. (H) Schematic of FDB single fiber culture experimental design. (I) Representative immunofluorescence (IF) images of KDR expression (yellow) in Pax7^+^ (red) satellite eYFP^-^ and eYFP^+^ (green) cells on FDB single myofibers isolated from wild type *Myf5-Cre:R26R-eYF*P mice. Nuclear DAPI (blue) stain was used. See also Figure S1F.

Satellite cells are cultured on single myofibers and fixed at 42h to visualize the first round of divisions, enabling the quantification of satellite stem cell symmetric and asymmetric divisions, as well as committed satellite cell divisions based on eYFP expression (Figure 1B). Positive controls for satellite stem cell asymmetric (EGF) and symmetric division (Wnt7a) were consistent with manual analysis, demonstrating a 1.3-fold change in divisions (Figure S1A and S1B). As previously described, our platform adapts the use of *flexor digitorum brevis* (FDB) muscles which are more appropriate for screening due to their size and satellite cell density (Shefer and Yablonka-Reuveni, 2005; Wang et al., 2019). Our novel screening platform is able to quantify changes in both asymmetric and symmetric stem cell fate decisions because it is based on imaging rather than PCR detection of Cre-mediated deletions (Wang et al., 2019).

We screened the Ontario Institute for Cancer Research (OICR) kinase inhibitor library containing over 480 kinase inhibitors using Wnt7a and EGF as positive controls for stimulating symmetric and asymmetric divisions, respectively (Le Grand et al., 2009; Wang et al., 2019). To assess positive hits, we devised a scoring system to identify targets of asymmetric division: targets with multiple compound inhibitors and effective IC50 values closer to the screening concentration of 1μM were weighted more heavily than targets with fewer inhibitors or IC50 values further away from 1μM. (Figure 1D). Our screen identified several previously reported targets of asymmetric division such as EGFR, Aurka, and p38-α/β MAPKs, validating the reliability of our platform to quantify asymmetric stem cell division (Chang et al., 2018; Wang et al., 2019). Notably, we identified the receptor tyrosine kinase KDR as a positive regulator of muscle asymmetric division (Figure 1C-D).

### KDR is expressed in satellite cells

Given the scarcity of satellite cells and difficulty in maintaining stem cell identity in culture, we validated KDR expression by quantitative digital droplet PCR (ddPCR) of freshly sorted satellite cells (FSC) from uninjured hindlimb *tibialis anterior* (TA) and *gastrocnemius* (GA) muscle, and proliferating myogenic precursor cells (PMC) isolated from hindlimb TA and GA muscle at 3 days post cardiotoxin injection (DPI) using *Pax7-nGFP* mice (Sambasivan et al., 2009) in both wild type and crossed with the *mdx* strains (Figure 1E). We followed the same gating strategy as previously reported with the addition of positive markers β1-Integrin (CD29) and VCAM1 (CD106) to further enrich for the stem cell population (Choo et al., 2017; Kuang et al., 2007; Rozo et al., 2016). We confirmed the purity of our isolations by quantifying myogenic regulatory factor (MRF) expression (Figure S1C).

Our ddPCR results confirm the expression of *Kdr* within satellite cells with greater expression in FSC compared to PMC populations in both wild type and *mdx* (Figure 1F). These results corroborate our transcriptome data on satellite cells and others (Dumont et al., 2015; Micheli et al., 2020). KDR protein expression was confirmed by immunofluorescence on satellite cells cultured on myofibers for 24h (Figure 1H-I). Interestingly, eYFP-expressing satellite myogenic cells expressed higher levels of KDR relative to eYFP^-^ satellite stem cells. Expression of KDR mRNA and protein in primary myoblasts was below the limit of detection (not shown). We also confirmed the expression of the KDR ligand *Vegfa* in both FSC and PMC populations (Figure 1G).

### KDR signaling modulates stem cell asymmetric division

To validate the effect of KDR signaling on stem cell asymmetric division and exclude possible off-target effects from small molecule compounds, we quantified satellite stem cell divisions following siRNA knockdown of *KDR* on *extensor digitorum longus* (EDL) myofibers from wild type *Myf5-Cre:R26R-eYFP* mice (Figure 2A). We confirmed the siRNA knockdown of *Kdr* using pre-validated commercially available qPCR and ddPCR primers on mouse-derived endothelial cells and by immunofluorescence (Figure S2).

**Figure 2.**
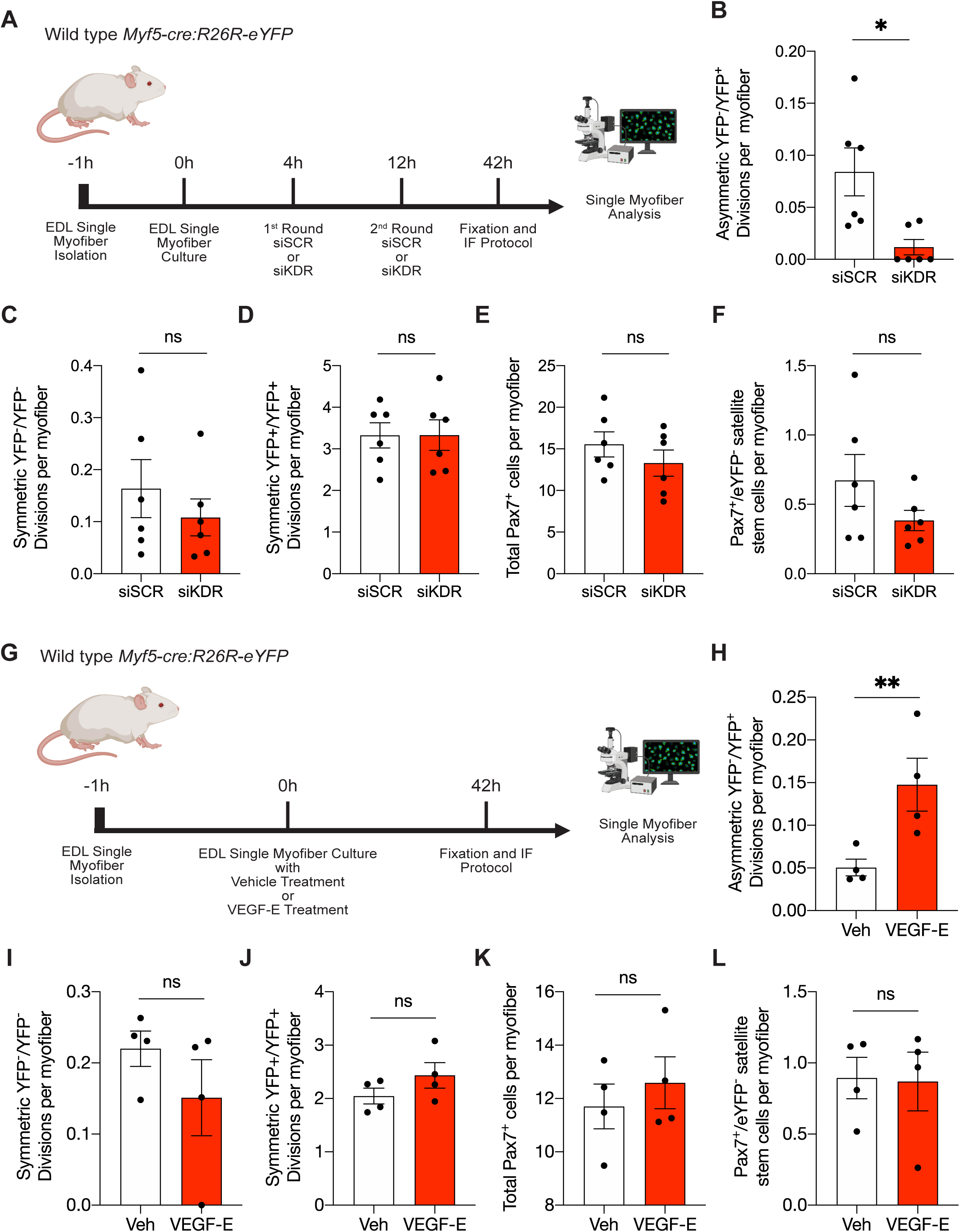
KDR modulates satellite stem cell asymmetric division. (A) Schematic of (B-F) siRNA knockdown experiments on EDL single myofiber isolated from wild type *Myf5-Cre:R26-eYFP* treated with control (siSCR, 10μM) or KDR knockdown (siKDR, 10μm). Analysis was performed after 42h of culture. Data is presented as mean. Error bars are ± SEM (n=6). *p<0.05. See also Figure S2A-F. (B) Numbers of Asymmetric YFP-/YFP+ divisions per myofiber (C) Numbers of Symmetric YFP-/YFP-divisions per myofiber. (D) Numbers of Symmetric YFP+/YFP+ divisions per myofiber. (E) Numbers of Pax7+ satellite cells per myofiber. (F) Numbers of Pax7+/eYFP-satellite stem cells per myofiber. (G) Schematic of (H-L) VEGFE ligand treatment experiments on EDL single myofiber isolated from wild type *Myf5-Cre:R26-eYFP* treated with vehicle (Veh) or VEGFE (100ng/ml). Analysis was performed after 42h of culture. Data is presented as mean. Error bars are ± SEM (n=4). *p<0.05 (H) Numbers of Asymmetric YFP-/YFP+ divisions per myofiber (I) Numbers of Symmetric YFP-/YFP-divisions per myofiber. (J) Numbers of Symmetric YFP+/YFP+ divisions per myofiber. (K) Numbers of Pax7+ satellite cells per myofiber. (L) Numbers of Pax7+/eYFP-satellite stem cells per myofiber.

Upon siRNA-mediated knockdown of *Kdr*, we observe a seven-fold decrease in satellite stem cell asymmetric divisions without a significant effect on the numbers of symmetric stem cell divisions (Figure 2B and 2C). This effect is specific to eYFP^-^ satellite stem cell divisions, as we observed no changes to numbers of eYFP^+^ satellite myogenic cell divisions (Figure 2D). Asymmetric stem cell divisions generate one stem (eYFP^-^) and one committed (eYFP^+^) daughter cell (Kuang et al., 2007). Consistent with this, we observed no change in the number of total Pax7^+^ satellite cells nor the number of eYFP^-^ satellite stem cells after the first round of divisions after 42h of culture (Figure 2E and 2F).

To further understand the role of KDR signaling in muscle stem cell asymmetric division, we explored the effects of activating the KDR signaling pathway. Given the promiuscuous binding of VEGF ligands and isomers to multiple VEGF receptors, we selected the use of Orf virus-derived VEGFE for its specificity in binding to KDR only (Ogawa et al., 1998; Vieira et al., 2010). VEGFE is able to bind to KDR homodimers and heterodimers with co-receptors Nrp1/Nrp2, with equal potency compared to VEGFA, without binding to decoy receptors VEGFR1 (also known as Flt1) or soluble VEGFR1 (sFlt1) (Figure 2G, S1D) (Kawamura et al., 2008; Ogawa et al., 1998; Vieira et al., 2010). Stimulation of the KDR signaling pathway with VEGFE ligand treatment increased the numbers of asymmetric divisions nearly 3-fold, corroborating the siRNA knockdown results (Figure 2H). Again, KDR signaling is specific to asymmetric stem cell division with no changes in symmetric satellite stem cell divisions, satellite myogenic cell divisions, total satellite cell numbers and satellite stem cell populations (Figure I-L).

Taken together, our data here suggests that KDR signaling specifically stimulates satellite stem cell asymmetric divisions. Interestingly, KDR signaling does not affect satellite stem cell fate decisions to undergo symmetric division, meaning satellite stem cell fate decisions are not necessarily binary.

### KDR signaling requires the dystrophin glycoprotein complex

Given the specificity of KDR signaling and its role in asymmetric but not symmetric division, we explored the role of KDR signaling in the context of DMD using *mdx* mice. Previously, we showed that satellite cells lacking dystrophin (*Dmd*) or dystroglyacn (*Dag1*) exhibit an inability to establish polarity leading to a significant reduction in numbers of asymmetric divisions and failure to generate sufficient numbers of committed progenitors for efficient muscle regeneration (Dumont et al., 2015).

To investigate KDR signaling in the DMD context, we isolated and cultured single EDL myofibers from *mdx* mice containing the *Myf5-Cre:R26R-eYFP* allele for quantification of satellite stem cell asymmetric divisions. Intriguingly, we observed no effect on asymmetric stem cell divisions following treatment of *mdx* single myofibers with VEGFE (Figure 3B). Moreover, VEGFE treatment of *mdx* myofibers did not affect satellite stem cell symmetric divisions, satellite myogenic cell divisions, or changes in satellite cell populations (Figure 3C-F).

**Figure 3.**
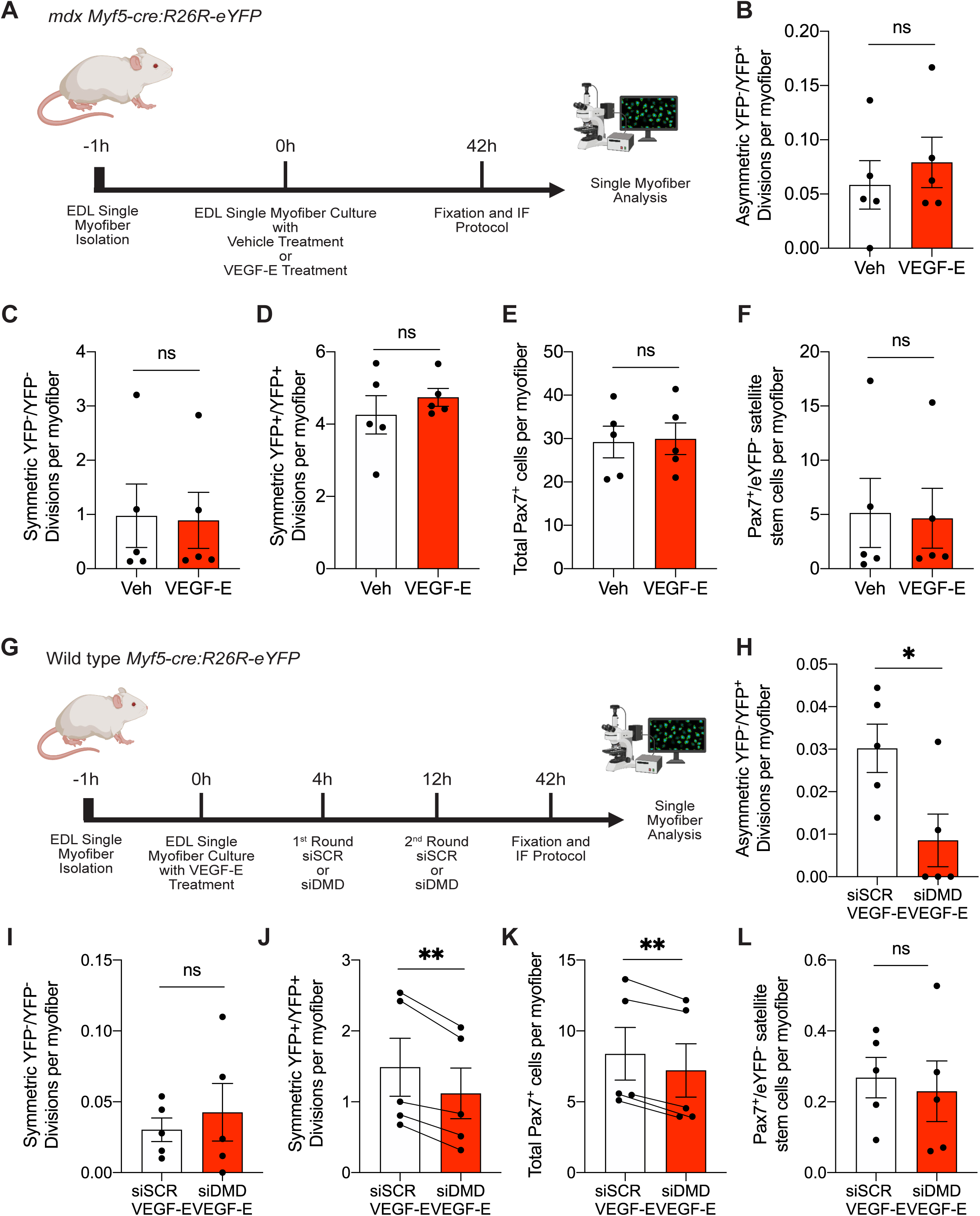
KDR signaling in MuSC requires the Dystrophin Glycoprotein Complex. (A) Schematic of (B-F) ligand treatment with vehicle (Veh) or VEGFE (100ng/ml) on EDL single myofiber isolated from *mdx Myf5-Cre:R26-eYFP*. Analysis was performed after 42h of culture. Data is presented as the mean. Error bars are ± SEM (n=5). (B) Numbers of Asymmetric YFP-/YFP+ divisions per myofiber (C) Numbers of Symmetric YFP-/YFP-divisions per myofiber. (D) Numbers of Symmetric YFP+/YFP+ divisions per myofiber. (E) Numbers of Pax7+ satellite cells per myofiber. (F) Numbers of Pax7+/eYFP-satellite stem cells per myofiber. (G) Schematic of (H-L) VEGFE (100ng/ml) treatment on control (siSCR, 10μM) or siDMD (10μM) treated EDL single myofibers isolated from wild type *Myf5-Cre:R26-eYFP*. Analysis was performed after 42h of culture. Data is presented as the mean. Error bars are ± SEM (n=5). *p <0.05, **p<0.001. See also Figure S3C-E. (H) Numbers of Asymmetric YFP-/YFP+ divisions per myofiber (I) Numbers of Symmetric YFP-/YFP-divisions per myofiber. (J) Numbers of Symmetric YFP+/YFP+ divisions per myofiber. (K) Numbers of Pax7+ satellite cells per myofiber. (L) Numbers of Pax7+/eYFP-satellite stem cells per myofiber.

To further investigate whether KDR signaling is overstimulated or impaired in *mdx* satellite cells, we performed siRNA-mediated knockdown of *Dmd* on single myofibers isolated from wild type *Myf5-Cre:R26R-eYFP* mice treated with VEGFE to mimic the *mdx* in an acute context (Figure 3G). The efficacy of siRNA knockdown of *Dmd* was validated by immunofluorescence and on wild type myoblasts after 2-days of differentiation by ddPCR using pre-validated commercially available *DMD* primers (Figure S3C-E).

VEGFE treatment was unable to rescue asymmetric divisions of satellite stem cells in conjunction with siRNA-mediated knockdown of *Dmd* (Figure 3H). In addition, no changes were observed in satellite stem cell symmetric divisions (Figure 3I). Consistent with previous findings in *mdx* mice, we observed fewer numbers of eYFP^+^ satellite myogenic cell divisions resulting in reduced numbers of total satellite cells following knock down of *Dmd* (Figure 3J and 3K) (Dumont et al., 2015). Moreover, we did not observe any change in numbers of eYFP^-^ satellite stem cells after 42h of culture (Figure 3L).

To investigate whether KDR interacts with the DGC complex, we performed a proximity ligand assay (PLA) on KDR and two components of the DGC complex: *Dmd* and *Dag1* in wild type satellite cells. Interestingly, we observed an increase in PLA signal with both components of the DGC complex, indicating close proximity between KDR and the DGC complex in satellite cells (Figure S3A and S3B). These data suggest KDR is closely associated with the DGC.

Our findings here show that the KDR-mediated satellite stem cell asymmetric division axis requires the DGC to function and the loss of DGC cannot be rescued by ligand stimulation of KDR.

### Reduced asymmetric divisions in KDR-deficient MuSC leads to fewer progenitors on cultured myofibers

Defects in asymmetric division can lead to cascading effects on the generation of progenitor cell populations. Following the first round of division around 42h, satellite cells rapidly undergo multiple rounds of division, forming clusters of satellite cells (Yin et al., 2013). To investigate these early effects following asymmetric division impairment, we crossed *Pax7^CreER^* (Murphy et al., 2011) with *KDR^flox/flox^* (Hooper et al., 2009) to delete *Kdr* specifically in Pax7-expressing satellite cells, hereafter referred to as KDR-KO. These mice were injected with tamoxifen for 5-days before EDL myofiber isolation and cultured with hydroxy tamoxifen (4-OHT). After culturing for 47h or 71h, myofibers were incubated with EdU one hour prior to fixation (Figure 4A). We utilized syndecan-4 (Syn4) rather than Pax7 to mark all satellite cells due to antibody species incompatibility with the anti-myogenin (MyoG) antibody.

**Figure 4.**
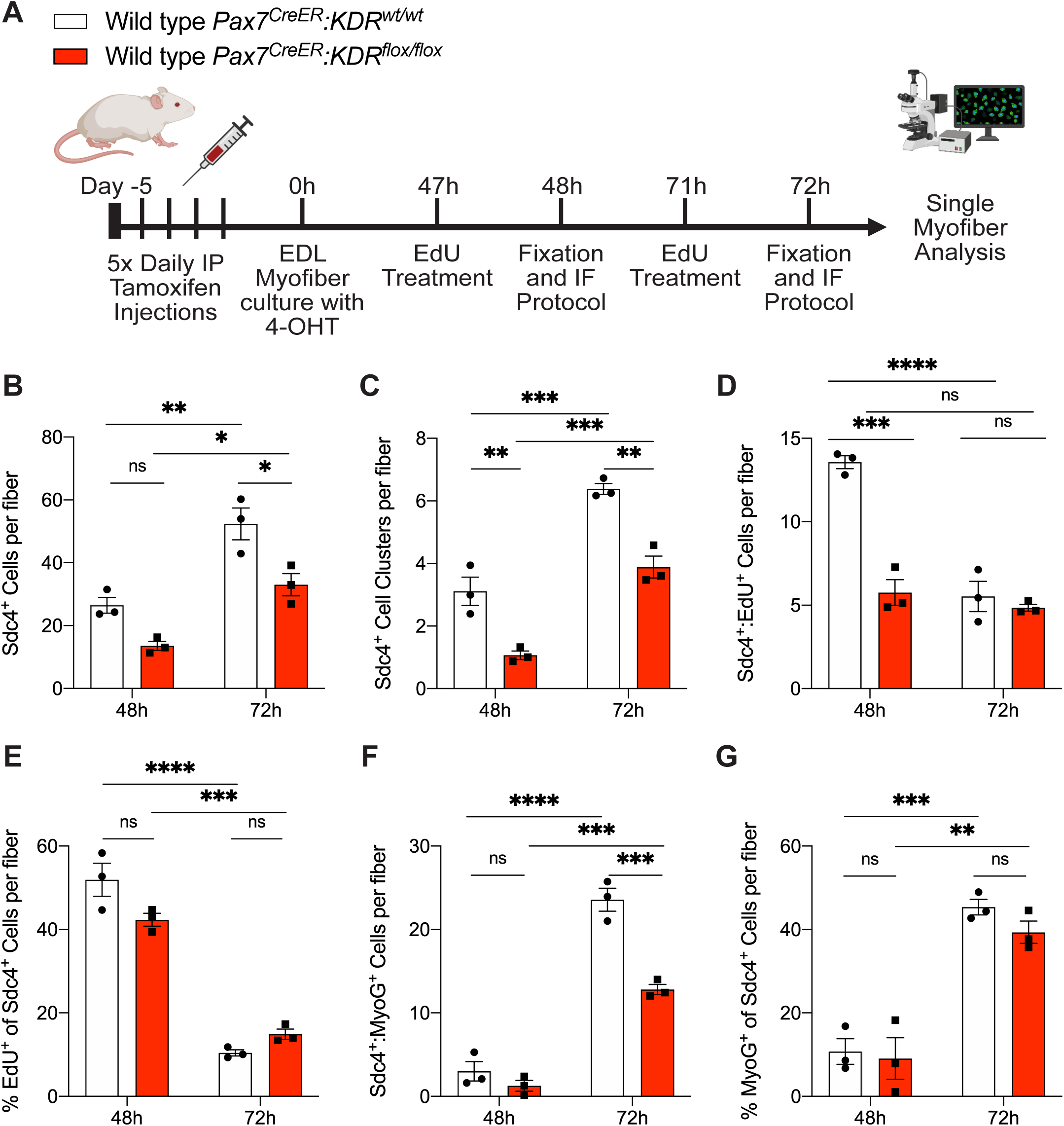
KDR-deficient MuSC generate reduced numbers of progenitors *ex vivo*. (A) Schematic of (B-I) EdU treatment on EDL single myofiber isolated from wild type *Pax7^CreER^:KDR^wt/wt^* (white) and wild type *Pax7^CreER^:KDR^flox/flox^* (red) mice cultured for 48h and 72h. Data is presented as mean. Error bars are ± SEM (n=3). *p<0.05, **p<0.01, ***p<0.001. (B) Total number of syndecan-4+ (Sdc4^+^) satellite cells per fiber. (C) Number of Sdc4^+^ satellite cell clusters (3+ cells). (D) Number of double positive Sdc4^+^:EdU^+^ cells per fiber. (E) Proportion of EdU^+^ cells of total Sdc4^+^ cells normalized per fiber. (F) Number of double positive Sdc4^+^:MyoG^+^ cells per fiber. (G) Proportion of MyoG^+^ cells of total Sdc4^+^ satellite cells normalized per fiber.

As expected, we observed a two-fold increase in satellite cell numbers at 72h compared to 48h in both wild type and KDR-KO fibers (Figure 4B). However, deficits in asymmetric division should be reflected in the population of early proliferating myogenic progenitors. Indeed, KDR-KO myofibers showed a 30% significant reduction in the numbers of Syn4^+^ satellite cells at 72h and a similar trend at 48h, indicating a reduced generation of early myogenic cells (Figure 4B). We also observed a marked reduction in the number of cell clusters containing >3 cells in close proximity at both 48h and 72h time points, corroborating downstream deficits in asymmetric stem cell division (Figure 4C).

Quantification of the number of dividing Syn4^+^EdU^+^ satellite cells revealed a 50% reduction in the numbers of dividing satellite cells at 48h in KDR-KO compared to wild type dividing satellite cells (Figure 4D). To determine whether the changes in Sdc4^+^ cell numbers were due to loss of asymmetric division or impaired proliferation, we then looked at the proportions of EdU^+^ actively dividing cells. We found no changes in total proportions of actively dividing EdU^+^ cells at both 48h and 72h time points suggesting KDR signaling is not involved in satellite cell proliferation kinetics (Figure 4E).

MyoG is expressed in myogenic cells that have withdrawn from the cell cycle and committed to terminal differentiation. Satellite cell derived progenitors expressing MyoG in KDR-KO cells were also reduced in number at 72h, with about half the number of MyoG-expressing cells present per fiber (Figure 4F). However, analysis of the proportions of MyoG^+^ cells relative to the total number of cells revealed no difference both at 48h and 72h (Figure 4G). Together, these data are consistent with the notion that the reduced levels of asymmetric division in the absence of KDR significantly reduce the generation of progenitors as reflected by the reduced numbers of MyoG-expressing progenitors undergoing commitment to differentiation.

Our results here provide further evidence that KDR signaling does not affect the rate of proliferation and differentiation of satellite cells. Rather, impairment of asymmetric division results in the reduced number of proliferative progenitors that are available to participate in repair.

### KDR-mediated asymmetric division is required for maintenance and repair

To investigate whether the effects of disrupting KDR-mediated asymmetric division persists throughout regeneration, we performed an *in vivo* injury experiment on KDR-KO and *Pax7^CreER^* mice. Cardiotoxin (CTX) injections were administered to injure one TA, while the undamaged TA in the contralateral leg was used as a control. The mice were allowed to recover for 10 days to enable quantification of dividing Ki67 satellite cells, MyoG^+^ progenitors, and fiber Feret’s minimal diameter (Figure 5A). After 10 days post injury (10-DPI), we found no difference in body weights or injured TA to body weight ratios (Figure 4SA and 4SB).

**Figure 5.**
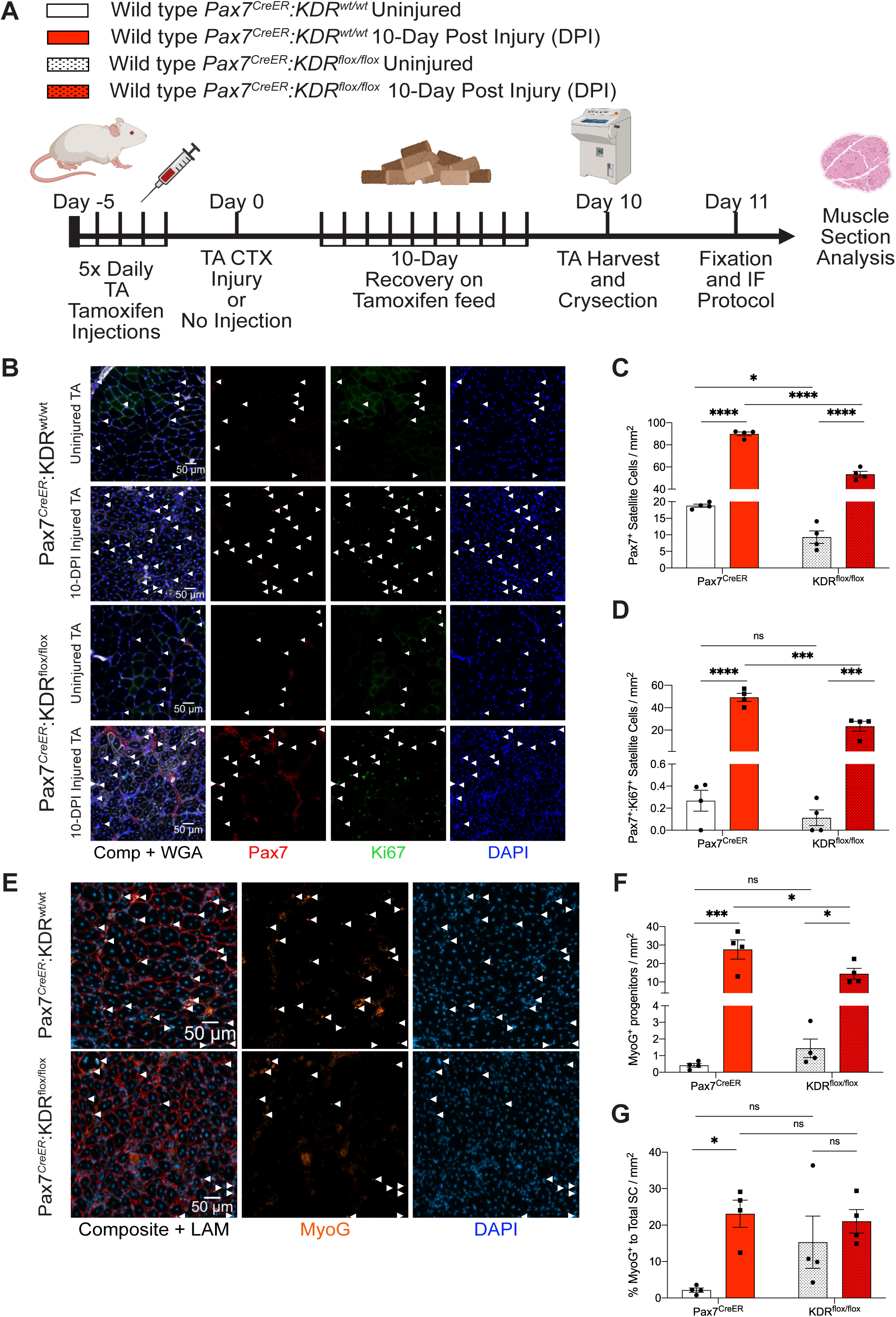
KDR-deficient MuSC generate reduced numbers of progenitors *in vivo*. (A) Schematic of (B-G) *Tibilais Anteri*o*r* (TA) Cardiotoxin (CTX) injury experiment on wild type *Pax7^CreER^:KDR^wt/wt^* and wild type *Pax7^CreER^:KDR^flox/flox^* mice. Data is presented as mean. Error bars are ± SEM (n=4). *p<0.05, **p<0.01, ***p<0.001, ****p<0.0001. (B) Representative immunofluorescence (IF) images of uninjured and injured TA muscle sections showing Pax7^+^ (red), Ki67^+^ (green), nuclei (DAPI) expression. Composite includes wheat germ (white). Arrows indicate Pax7^+^ satellite cells. See also Figure S5. (C) Quantification of Pax7^+^ satellite cell numbers normalized to area in uninjured vs injured TA sections from *Pax7^CreER^:KDR^wt/wt^* and wild type *Pax7^CreER^:KDR^flox/flox^* mice. (D) Quantification of Pax7^+^Ki67^+^ satellite cell numbers normalized to area in uninjured vs injured TA sections from wild type *Pax7^CreER^:KDR^wt/wt^* and wild type *Pax7^CreER^:KDR^flox/flox^* mice. See also Figure S54D. (E) Representative immunofluorescence (IF) images of uninjured and injured TA muscle sections showing Pax7^+^ (red), MyoG^+^ (orange), nuclei (DAPI) expression. Composite includes Laminin (red). Arrows indicate MyoG^+^ cells. See also Figure S6. (F) Quantification of MyoG^+^ cell numbers normalized to area in uninjured vs injured TA sections from wild type *Pax7^CreER^:KDR^wt/wt^* and wild type *Pax7^CreER^:KDR^flox/flox^* mice (G) Proportion of Pax7^+^ satellite cells that are MyoG^+^ normalized to area in uninjured vs injured TA serial sections from wild type *Pax7^CreER^:KDR^wt/wt^* and wild type *Pax7^CreER:^KDR^flox/flox^* mice.

Interestingly, we observed that KDR deletion resulted in a significant 2-fold decrease in satellite cell numbers that corresponded with a trend of increased MyoG progenitors and centrally nucleated fibers in uninjured KDR-KO TA muscles at 10 days, after tamoxifen treatment suggesting a propensity for fusion (Figure 5B, 5C, 5F, S5, 6B, and S6). At 10 days following cardiotoxin induced injury, we observed a 1.6-fold decrease in numbers of Pax7^+^ satellite cells in the TA from KDR-KO mice compared to *Pax7^CreER^* control counterparts (Figure 5C). Consequently, we also observed a reduction in the numbers of actively dividing Ki67^+^ satellite cells (Figure 5D and S4E).

To investigate whether the reduction in satellite cell numbers is due to reduced proliferation or a reduced early progenitor population from impaired asymmetric division, we compared both the proportion of Ki67^+^ satellite cells and population change in response to injury and found no difference between KDR-KO and *Pax7^CreER^* control mice (Figure S4C and S4D). These findings indicate the rate of proliferation in both KDR-KO and wild type are comparable, and the deficits seen in satellite cell numbers 10 days post injury are due to reduced generation of proliferating progenitors as a consequence of the reduction in asymmetric satellite stem cell divisions.

To investigate differentiation kinetics, we quantified the numbers of myogenic cells expressing MyoG in both KDR-KO and *Pax7^CreER^* control mice (Figure 5E, 5F and S6). As expected, MyoG^+^ satellite cells increased dramatically at 10 days following injury (Figure 5F). In uninjured TA muscle, we observed no significant difference in the numbers of MyoG-expressing cells when comparing KDR-KO to *Pax7^CreER^* control mice (Figure 5F). In line with our previous results on KDR-KO myofibers, we observed a 2-fold decrease in the numbers of MyoG^+^ cells in injured KDR-KO compared to *Pax7^CreER^* control TA (Figure 4F and 5F). Furthermore, KDR signaling does not appear to affect differentiation kinetics even at 10 days post injury despite fewer MyoG^+^ cells, as we observed no difference when comparing the proportion of MyoG^+^ cells in injured TA of KDR-KO and *Pax7^CreER^* mice (Figure 5G). These findings are consistent with our earlier observations that KDR does not affect the rate of differentiation, and that deficits seen in progenitor populations stem from a reduced number of asymmetric satellite stem cell divisions.

To determine the impact of the loss of KDR and asymmetric division on muscle regeneration, we analyzed the number and size of myofibers (Figure 6). We observed no significant difference in the total number of fibers or centrally nucleated fibers in KDR-KO TA muscle (Figure 6A and 6B). However, quantification of Feret’s fiber diameter revealed a significant increase in the numbers of smaller myofibers (Figure 6C, 6D). Indeed, the distribution of the Feret’s fiber diameter shows a significant leftward shift indicating the increase in numbers of smaller sized fibers (Figure 6E and 6F).

**Figure 6.**
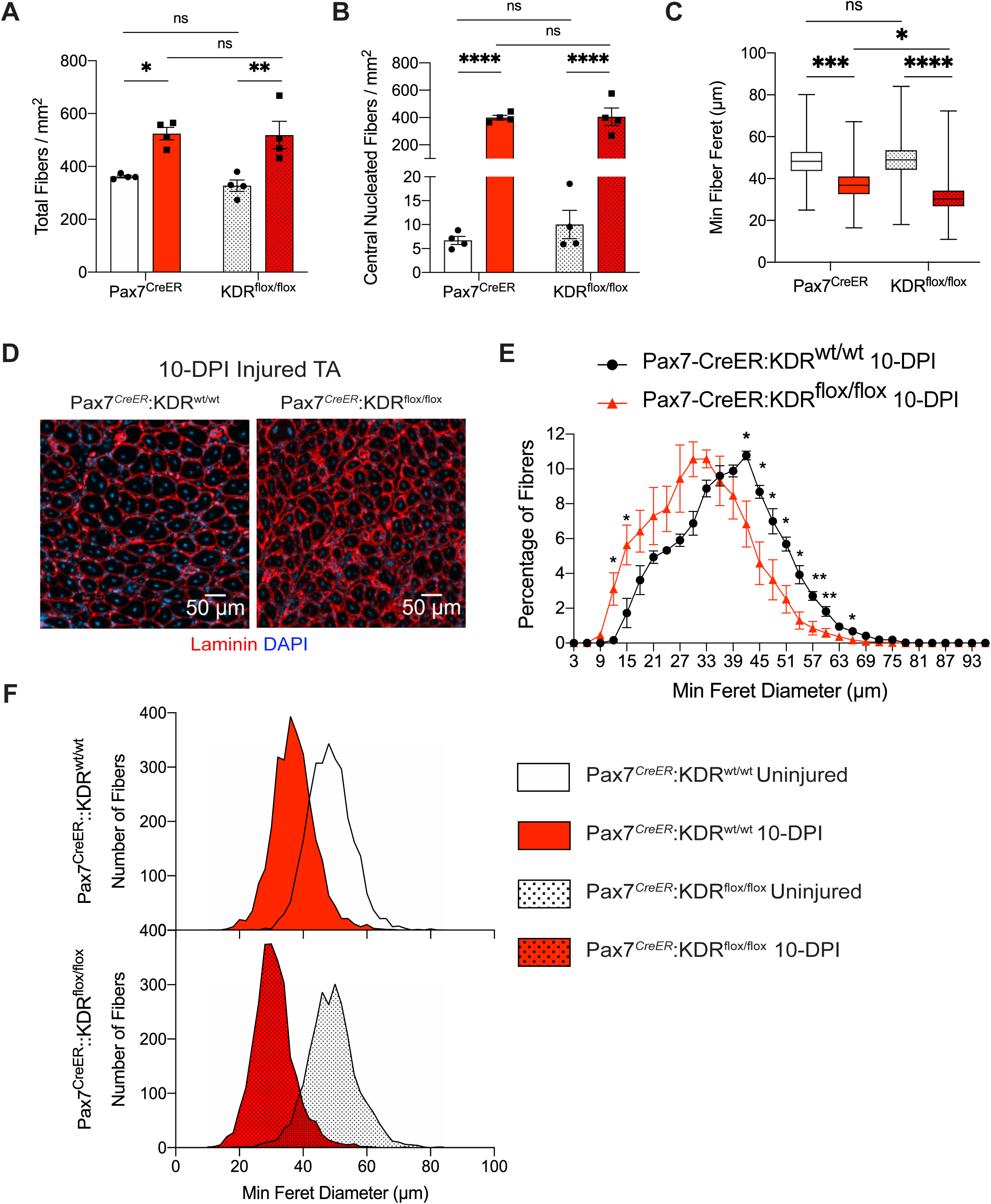
Impaired muscle regeneration following KDR-deletion in MuSC. (A) Quantification of fiber numbers normalized to area in uninjured vs injured TA from wild type *Pax7^CreER^:KDR^wt/wt^* and wild type *Pax7^CreER^:KDR^flox/flox^* mice. Data is presented as mean. Error bars are ± SEM (n=4). *p<0.05, **p<0.01. See also Figure S4E. (B) Quantification of central nucleated fibers normalized to area in uninjured vs injured TA from wild type *Pax7^CreER^:KDR^wt/wt^* and wild type *Pax7^CreER^:KDR^flox/flox^* mice. Data is presented as mean. Error bars are ± SEM (n=4). ****p<0.0001. See also Figure S4F. (C) Mean comparisons of fiber ferret diameter (μm) in uninjured vs injured TA from wild type *Pax7^CreER^:KDR^wt/wt^* and wild type *Pax7^CreER^:KDR^flox/flox^* mice. Data is presented as mean. (n=4). *p<0.05, ***p<0.001, ****p<0.0001. See also Figure S4G. (D) Representative immunofluorescence (IF) images of 10-Day post injured (10-DPI) TA muscle sections from wild type *Pax7^CreER^:KDR^wt/wt^* and wild type *Pax7^CreER^:KDR^flox/flox^* mice stained with Laminin (red) and DAPI (blue). See also Figure S6. (E) Histogram overlay of minimum fiber ferret diameter (μm) comparing 10-DPI wild type *Pax7^CreER^:KDR^wt/wt^* vs wild type *Pax7^CreER^:KDR^flox/flox^.* Data is presented as mean. Error bars are ± SEM (n=4). *p<0.05. (F) Distribution of minimum fiber ferret diameter (μm) comparing uninjured vs injured TA from wild type *Pax7^CreER^:KDR^wt/wt^* and wild type *Pax7^CreER^:KDR^flox/flox^* mice. See also Figure S4H.

Our evidence demonstrates that KDR-signaling is required for stimulation of asymmetric division and its loss handicaps the early generation of committed satellite cells required for the exponential expansion of myogenic progenitors for repair. However, KDR-signaling does not affect the downstream rate of myogenic proliferation and differentiation during regeneration. This cascading effect on regeneration demonstrates the importance of asymmetric division in muscle repair as well as maintenance.

## DISCUSSION

This is the first report of KDR-mediated signaling and its physiological role in muscle satellite stem cells. In this study, we show how KDR signaling is a critical component of stem cell fate decisions to undergo asymmetric division and that it requires a polarized intrinsic cell environment established by the DGC complex (Dumont et al., 2015). KDR signaling is specific to asymmetric division and its manipulation does not affect satellite stem cell symmetric division nor downstream rate of progenitor cell proliferation and differentiation. Notably, we observe a 2-fold decline in the numbers of satellite cells following tamoxifen treatment suggesting that KDR signaling may also have a role in preventing precocious differentiation of satellite cells in quiescence. Our study provides important new insight into the regulation of stem cell asymmetric divisions in muscle maintenance, repair, and disease states.

The prevailing theory on stem cell fate is that asymmetric versus symmetric division is a binary decision. Indeed, past investigations on cell polarity pathways have demonstrated increased stem cell asymmetric or symmetric division when the other is perturbed (Bentzinger et al., 2013; Kuang et al., 2007; Wang et al., 2019). This study provides the first evidence of a signaling pathway that exclusively affects asymmetric division without impacting symmetric division (Figure 2B, 2C, 2H, 2I). This finding is a significant advance in our understanding of the role of asymmetric division in disease and aging (Bernet et al., 2014; Dumont et al., 2015).

Here we identify KDR and possible co-receptors Nrp1/Nrp2 expression in satellite stem cells (Figure 1F, 1G,1I and S1D). Asymmetric division occurs during the first few rounds of division (Feige et al., 2018; Kuang et al., 2007; Yin et al., 2013). Accordingly, KDR expression is correlated with asymmetric stem cell division, and its expression is strongly downregulated in actively proliferating committed satellite cells (Figure 1F). VEGFR1 (Flt-1), a known decoy receptor that competitively and preferentially binds to VEGFA ligand over KDR (Tanaka et al., 1997), also demonstrated a similar expression profile to KDR and was downregulated in proliferating committed satellite cells (Figure S1D). Intriguingly, we did not detect Flt-1 expression by ddPCR from sorted *mdx* satellite cells (Figure S1D). Downregulation of Flt-1 in *mdx* satellite cells may be due to an attempt to compensate for impaired KDR signaling by the removal of VEGFR1-mediated negative feedback on KDR signaling.

Our gene expression analysis indicates that VEGFA is the primary ligand expressed in satellite cells (Figure S1E). Other sources of VEGFA in the muscle include myofibers, endothelial cells, and FAPs (Blackburn et al., 2019; Verma et al., 2018). The KDR signaling pathway is tightly controlled with numerous negative feedback loops to prevent overactivation (Simons et al., 2016). Given the abundant availability and redundant expression of VEGFA from multiple sources, it is likely that KDR-mediated asymmetric signaling occurs primarily in an autocrine fashion, or short-range paracrine signaling (i.e. myofiber-satellite cell signaling). Indeed, VEGFA knockout in MuSC or FAPs demonstrate different regenerative outcomes on myofiber size and capillary density respectively, suggesting VEGFA is a short-range ligand that does not easily pass through the basal lamina to affect other cell types (accompanying paper in this issue Groppa et al., 2020).

Dystrophin, a critical component of the DGC, is required for establishing cell polarity in muscle stem cells, important for asymmetric division (Dumont et al., 2015). Our findings show that KDR is closely associated with the DGC complex, and that impaired KDR-signaling in DMD cannot be rescued by ligand treatment, suggesting KDR signaling occurs downstream of DGC formation (Figure 3, S3). However, whether KDR interacts directly or indirectly with the DGC in modulation of stem cell asymmetric division remains to be explored.

Given the complexity of receptor tyrosine kinase activity and cross talk potential, KDR signaling may be interacting with other asymmetric stem cell signaling pathways. Indeed, studies in cancer biology have demonstrated VEGF promotes self-renewal in cancer stem cells through KDR/STAT3 signaling, and stimulate phosphorylation of p38α MAPK, two known pathways of muscle stem cell asymmetric division (Bernet et al., 2014; Masson-Gadais et al., 2003; Nilsson et al., 2016; Price et al., 2014; Saulle et al., 2009; Troy et al., 2012; Zhao et al., 2015).

Our *ex vivo* and *in vivo* observations on KDR-KO mice suggest that an impairment of asymmetric division leads to a decline in the numbers of early myogenic progenitors, with no effect on proliferation and differentiation kinetics (Figure 4, 5, S4, S5 and S6). Analysis of KDR-KO on non-injured TA muscles revealed a 50% decline in satellite cell numbers 10 days following tamoxifen treatment, suggesting the onset in satellite cell number decline begins during steady state (Figure 5C). Corroborating with our findings of reduced satellite cell numbers in non-injured TA muscle, we observed that KDR-KO exhibited a trend of increased MyoG numbers, proportion of MyoG-expression satellite cells and centrally nucleated fibers that (Figure 5F, 5G, 6B and 5C). These observations suggest that satellite cell specific loss of KDR in uninjured muscle promotes premature fusion through a break in quiescence.

The mechanism of action behind this decline may be associated with the chemoattractant functions of VEGFA signaling. Satellite stem cell derived VEGFA secretion has recently been reported to recruit endothelial cells, required for maintaining stem cell quiescence via Notch signaling (Verma et al., 2018). Similarly, myofiber secretion of VEGFA under homeostatic conditions may act as a chemoattractant to maintain quiescence by securing satellite cells within their niche (Baghdadi et al., 2018a; Blackburn et al., 2019). Observations on satellite cell numbers following satellite cell specific VEGFA-KO complement our findings in KDR-KO mice, and demonstrate a similar loss in early myogenic numbers (accompanying paper in this issue Groppa et al., 2020). The similarities between KDR-KO and *mdx* phenotype include impaired asymmetric division and precocious activation of satellite cells. However, the *mdx* phenotype exhibits stem cell hyperplasia not present in KDR-KO, and cell division defects that are not limited to satellite asymmetric stem cell division (Dumont et al., 2015). The similarities in muscle phenotype between KDR-KO and *mdx* reinforces the importance of asymmetric stem cell division in muscle repair, while the differences suggests other compensatory mechanisms are at play in the context of chronic injury.

Our results provide compelling evidence that KDR-mediated signaling is restricted to satellite stem cell fate decision and does not impact downstream proliferation or differentiation kinetics of progenitors (Figure 4 and Figure 5). These findings suggest that the role of asymmetric division in response to acute injury is to generate sufficient numbers of satellite myogenic cells that are then able to undergo multiple but limited rounds of division. In accordance with our results, muscle satellite cell specific VEGFA knockout mice show reduced numbers of Pax7-expressing cells up to 12 weeks post injury (Verma et al., 2018). Thus, the marked reduction in satellite stem cell asymmetric divisions results in the generation of insufficient numbers of proliferating myogenic progenitors to efficiently undertake the regeneration of muscle (Figure 5 and Figure 6).

Our results show that KDR signaling is critical for asymmetric cell division and requires the establishment of the DGC complex to function and suggest that this signaling pathway may be perturbed in DMD patients. Furthermore, KDR-mediated stimulation of asymmetric division is important for both maintaining satellite cell numbers, and the initial generation of proliferating committed cells that is required for efficient muscle repair in response to acute injury. Future studies will elucidate the role of KDR signaling in larger context of stem cell polarity and the potential for small molecule activation of KDR as a therapeutic modality for enhancing muscle repair.

## STAR METHODS

### Key Resources Table

See document Key Resource Table.

### Contact for Reagent and Resource Sharing

Information and requests for reagents may be directed to the Lead Contact, Michael A. Rudnicki.

### Experimental Model and Subject Details

#### Experimental animals

We performed all experiments in accordance with the Canadian Council Animal Care Guidelines and the University of Ottawa Animal Care Committee. We used adult 2-3 month old mice for all experiments. Mice are housed in ventilated cages and ¼” corncob bedding is changed every 14 days. Mice are supplied food *ad libitum* with automated acidified RO water or bottles changed every 7 days. Light cycles are 12-12h cycle with simulated sunrise and sunset. Health and immune status of experimental mice were normal where Sentinels are routinely monitored (fecal, fur, and oral swab) for *MNV, MHV,* Mouse *Pneumocystis*, pinworm, and *Spironucleus muris*. No animals were subjected to previous experimental procedures.

The following mouse lines were used: C57BL/B10ScSn-*Dmd*^mdx^/J (mdx, homozygous *Dmd*^mdx^ females and hemizygous *Dmd*^mdx^ males). B6.Cg-*Pax7*^tm1(cre/ERT2)Gaka^/J (*Pax7^CreER^*), B6.129S4*-Myf5^tm3(cre)Sor^*/J (*Myf5-Cre*), B6.129X1-*Gt*(*ROSA*)*26Sor^tm1(EYFP)Cos^*/J(*R26R-eYFP*), B6.SJL-*Pax7^tm1.2Tajb^* (*Pax7-nGFP*) (Bulfield et al., 1984; Murphy et al., 2011; Sambasivan et al., 2009; Srinivas et al., 2001; Tallquist et al., 2000).

*Myf5-Cre:ROSA26-eYFP* (Kuang et al., 2007) mice were F1 progeny from *Myf5-Cre* x *R26R-eYFP* breeding pairs. Tamoxifen inducible conditional genetic knockout animals were F2 crosses between offspring of *Pax7-Cre^ER/+^* mice and *KDR^fl/fl^* (Hooper et al., 2009) mice generating Pax7-Cre^ER/+^:KDR*^+/+^* and Pax7-Cre^ER/+^:KDR*^fl/fl^*. Unless otherwise stated, 6-8 week old mice were used for all experiments.

#### Primary cell acquisition and culture

Primary endothelial cells were isolated and cultured as described (Ouellette et al., 2020). Cortical tissue was dissected, minced, and incubated in Neural Tissue Dissociation compounds (Miltenyi Biotec). Primary endothelial cells were then isolated using CD31 microbeads (Miltenyi Biotec) before cultured in an EC culture medium (PromoCell). Culture medium was replaced 24h post-seeding and every 48h until confluency. Mouse-derived primary endothelial cells were used for validation of KDR expression following siRNA knockdown.

Depending on the purity required, satellite cells were isolated either by FACS or MACS. FACS isolation was performed as previously described using *Pax-nGFP* mcie (Wang et al., 2019). For all other acquisition of satellite cell lines, MACs isolation was performed as previously described (Chang et al., 2018). Mouse-derived myoblasts (Passage <10) were used for validation of DMD siRNA knockdown. These cells were isolated using the MACS Satellite Cell Isolation Kit (Miltenyi Biotec) and anti-α7-integrin beads (Miltenyi Biotec). Primary myoblasts were cultured in Ham’s F10 supplemented with 20% FBS, 2% Chicken embryo extract, 1% penicillin/streptomycin and 2.5 ng/ml bFGF. Myoblasts were differentiated for 2 days in DMEM supplemented with 10% horse serum, 1% penicillin/streptomycin and 2.5 ng/ml bFGF and treated with siRNA before mRNA was collected.

#### Lineage tracing with Myf5Cre:R26R-eYFP and Pax7-nGFP

*Pax7-nGFP* transgenic mice possess a knock-in of nuclear localized EGFP (nGFP) within the first exon of Pax7(Sambasivan et al., 2009). Expression of nGFP corresponds with endogenous Pax7 gene expression. *Pax7-nGFP* mice allow for the visualization of Pax7-expression satellite cells. Myf5-Cre:R26R-eYFP transgenic mice possess a knock-in of Cre recombinase within the 5’-UTR of the Myf5 gene crossed with a knock-in of Cre-mediated activation of yellow fluorescent protein (eYFP) within the Gt(ROSA)26Sor locus(Srinivas et al., 2001). Myf5-Cre:R26R-eYFP mice allow for the differentiation between committed satellite cells that have expressed Myf5-Cre and satellite stem cells that have never expressed Myf5-Cre (Kuang et al., 2007).

#### Small molecule compound library

The Ontario Institute for Cancer Research kinase inhibitor library (OICR, Toronto, Canada) consists of 400 specific kinase inhibitors and 160 additional small molecule compounds targeting cellular and development pathways. Compound libraries were obtained as 1mM solutions dissolved in DMSO.

#### Single myofiber culture and siRNA transfection

Single myofibers were isolated and cultured as previously described (Dumont et al., 2015; Wang et al., 2019). EDL or FDB muscles were dissected and incubated at 37°C in DMEM with 2% L-glutamine, 4.5% glucose and 110mg/mL sodium pyruvate (Gibco) containing 0.2% collagenase I (Sigma) for 1h (EDL) or 3h (FDB). Myofibers were separated by trituration using a glass pipette. Myofibers were cultured at 37°C in DMEM with % L-glutamine, 4.5% glucose and 110mg/mL sodium pyruvate (Gibco) supplemented with 20% FBS (Wisent), 1% chicken embryo extract (MP Biomedicals) and 2.5 ng/ml bFGF (Cedarlane). FDB screening assay culture media was supplemented with DMSO (1:1000 dilution; Sigma), DMSO + EGF (100ng/mL; Miltenyi Biotec), DMSO + Wnt7a (50 ng/mL; R&D Systems), or 1 μM of small molecule compound. Fibers cultured for 42h before fixation and immunofluorescence work up. Fibers were imaged on the Cellomics High Content platform and satellite stem divisions were quantified on a custom MATLAB-based script using the Cellomics output data.

(Dumont et al., 2015; Wang et al., 2019)For EGF treatments, human recombinant EGF (100ug/ml in PBS with 0.1% BSA) was added to culture medium at 100 ng/ml with 1% BSA in PBS as vehicle control. For Wnt7a treatments, human recombinant Wnt7a (50ug/ml in PBS with 0.1% BSA) was added to culture medium at 50 ng/ml with 1% BSA in PBS as vehicle control. For VEGFE treatments, ORF viral VEGFE (100ug/ml in PBS with 0.1% BSA) was added to culture medium at 100 ng/ml with 1% BSA in PBS as vehicle control. For EdU labeling, EdU was supplemented into media 1h prior fixation as per manufacturer’s protocol.

Transfection of satellite cells on myofibers was performed using the lipofectamine RNAimax (Life Technologies) and validated smartpool siRNAs for *KDR*,*DMD*, or scramble (SCR) (Integrated Technologies). Two transfections were performed at 4h and 16h post isolation of myofibers as previously described (Le Grand et al., 2009). (Table S1). (Wang et al., 2019)

#### Cardiotoxin and tamoxifeni Injections

For TA cardiotoxin (CTX) injuries, intramuscular cardiotoxin injections (Latoxan, 50μl of 10 μM solution in saline) were injected into TA muscle through skin under general anesthesia. For hindlimb injuries, two additional CTX injections were made to the thigh and gastrocnemius muscle. Tamoxifen (cat#T5648, Sigma) was diluted in corn oil (20mg/mL). Intraperitoneal injections were administered at a dosed of 100ul/ 20g mouse.

#### Immunofluorescence and immunohistochemistry

EDL myofibers were fixed with 2% PFA for 10 min. Fibers were blocked and permeabilized (5% horse serum, 1% BSA (Sigma), 0.5% Triton X-100 (Sigma), PBS) for 1h at room temperature. Primary antibody diluted in blocking solution was then incubated either overnight at 4°C or room temperature for 2h. Samples were then incubated with conjugated Alexa Fluor (1:1000, Life Technologies) secondary antibodies diluted in blocking solution for 1h. Nuclei were counterstained with DAPI (D8417, Sigma) in PBS for 10 min and mounted with PermaFluor (Fisher). Antibodies used include: mouse anti-Pax7 (DSHB), chicken anti-GFP (cat# ab13970, abcam), goat anti-KDR (cat# AF644 R&D Systems), rabbit anti-KDR (cat# ab39256, abcam), rabbit anti-DMD (cat# ab15277, abcam), rabbit anti-Dag1 (cat# D1945, Sigma), and chicken anti-syndecan-4 (Cornelison et al., 2004).

Tibilialis Anterior (TA) muscle cryosections were fixed with 2%PFA for 10 min and permeabilized (0.1% Triton X-100, 0.1M Glycine, PBS) for 5 min. Muscle sections were washed with PBS and blocked with M.O.M blocking reagent (Vector Laboratories) for 2h. Primary antibody incubations were performed at 4°C overnight. TA sections were then incubated with conjugated Alexa Fluor (1:1000, Life Technologies) secondary antibodies and wheat germ agglutinin (W32466, Life Technologies) diluted in blocking solution for 1h. Sections were then stained with DAPI and mounted with PermaFluor mounting medium (Fisher). Antibodies used include: mouse anti-Pax7 (DSHB), rabbit anti-Ki67 (cat# ab39256, abcam), mouse anti-MyoG (cat# MAB66861, R&D Systems), and rat anti-laminin (cat# L0663, Sigma).

#### Histological analysis of muscle sections

Central nucleated fibers were quantified manually in relation to Wheat Germ Agglutinin fluorescence signal. Pax7, Ki67, and MyoG cells were quantifications were automated using the IMARIS software (Oxford Instruments) and validated manually. Laminin staining was used to quantify minimum Fiber diameter. Minimum fiber ferret diameter measurement was performed using SMASH standalone software as described previously (Smith and Barton, 2014). Total number of fibers was manually validated.

#### Proximity Ligation Assay (PLA)

Proximity ligation assays were performed on fixed myofibers as previously described (Chang et al., 2018; Dumont et al., 2015; Wang et al., 2019). Myofibers were permeabilized (0.1% Trition X-100, 0.1M Glycine, PBS) for 10 min and blocked with Duolink Blocking Solution (Sigma) for 2h. Myofibers were incubated overnight at 4°C with primary antibodies diluted in Duolink Blocking Solution (Sigma). Duolink PLA probes (Sigma) rabbit and goat were used and Duolink *In Situ* Detection Reagents Red (Sigma) were performed as per manufacturer’s instructions. Images of were acquired on an Axio Observer.Z1 microscope equipped with a LSM510 META confocal laser scanner and a plan-Apochromat 63x/1.40 Oil DIC M27 objective or an Axioplan 2 microscope equipped with a plan-Neofluar 40x/1.30 Oil DIC and a plan-Neofluar 100x/1.30 Oil DIC objective. Images were processed and analyzed with Axiovision, Zen, and FIJI software. 3D z-stack images were projected by maximum intensity using Fiji software **Error! Hyperlink reference not valid.**p**Error! Hyperlink reference not valid.** Quantification of PLA was performed by counting PLA puncta per satellite cell.

#### Real-Time PCR

RNA extractions were performed using Trizol and the Nucleospin RNA II kit (Macherey-Nagel) as per manufacturer’s instructions. cDNA reverse transcription was performed using the iScript cDNA synthesis kit (Bio-Rad). SYBR Green, real-time PCR analysis (SSoFast EvaGreen Supermix; BioRad) was performed on the CFX384 real time PCR detection system (Bio-Rad) and normalized to *PPIA*, *RSP18* and *GAPDH*. Results were analyzed by Bio-Rad CFX Manager Software (Table S2).

#### Digital droplet PCR

RNA extraction on sorted satellite cells was performed using the Arcturus Picopure RNA extraction kit (Thermo Fisher Scientific) and cDNA reverse transcription was performed using the iScript cDNA synthesis kit (Bio-Rad) according to manufacturer’s instructions. ddPCR was performed using ddPCR Supermix for Probes (no dUTP) (Bio-Rad) according to manufacturer’s instructions and analyzed with QX100 Droplet Digital PCR System (Bio-Rad). Quantitative mRNA copies were normalized to *TBP*. Probes were purchased from Integrated DNA Technologies (Table S2).

#### Quantification and statistical analysis

Compiled data are expressed as mean ± standard error of the mean (SEM). Experiments were performed with a minimum of three biological replicates unless otherwise stated. For statistical comparisons of two conditions, Student’s t-test was used. Paired tests were used when comparing biologically matched samples. Unpaired tests were used when comparing biologically unrelated samples. Ferret size measurements were analyzed using one-way non-pairing ANOVA with Tukey’s multiple comparisons test. No data were removed as outliers with graphic data point representation included where appropriate. Experimental design incorporated user blinding when possible. Statistical analysis was performed in GraphPad Prism and Microsoft Excel. The level of significance is indicated as follows: *p<0.05, **p<0.01, ***p<0.001, ****p<0.0001.

#### Resource availability

The datasets supporting the current study are available from the corresponding author on request.

## SUPPLEMENTAL INFORMATION

Supplemental information includes five figures.

## ACKNOWLEDGMENTS

The OICR kinase inhibitor library was kindly provided by Dr. Rima Al-Awar. We thank Dr. Julian Yokell-Lelievre for support and guidance during the development of our high content screen. We thank David Cook for his help with MATLAB visualizations of our screen results. We thank Dr. Natasha Chang, Dr. Caroline Brun, and David Datzkiw for their guidance with our ddPCR, EdU, and PLA studies. We thank Julie Ouellette and Dr. Baptiste Lacoste for their guidance in isolating mouse endothelial cells. We thank Fernando Ortiz and the Cell Sorting and Flow Cytometry Core Facility at the OHRI. Graphical abstract was created with BioRender.com. W.C. is supported by fellowships from QEII-GSST and University of Ottawa Brain and Mind Research Institute Éric Poulin Centre for Neuromuscular Disease Scholarship in Translation Research Award. M.A.R. holds the Canada Research Chair in Molecular Genetics. The studies from the laboratory of M.A.R. were carried out with support of grants from the US National Institutes for Health [R01AR044031], Canadian Institutes for Health Research [FDN-148387], and the Stem Cell Network.

## AUTHOR CONTRIBUTIONS

Conceptualization, W.C. and M.A.R.; Methodology, W.C., Y.X.W., T.J.P, M.R., and M.A.R.; Investigation, W.C., M.R., M.A.R.; Writing – Original Draft, W.C. and M.A.R.; Writing – Review & Editing, W.C., Y.X.W., M.R., D.D., A.L., J.S., T.J.P, M.R., M.A.R.; Funding Acquisition, M.A.R.; Resources, M.A.R; Supervision, M.A.R.

