## Supplemental Data for "KDR Signaling in Muscle Stem Cells Promotes Asymmetric Division and Progenitor Generation for Efficient Regeneration"

1. Sprott Center for Stem Cell Research
   Ottawa Hospital Research Institute
   Regenerative Medicine Program
   Ottawa, ON, Canada

2. Department of Cellular and Molecular Medicine
   Faculty of Medicine
   University of Ottawa
   Ottawa, ON, Canada

3. Department of Medicine
   Faculty of Medicine
   University of Ottawa
   Ottawa, ON, Canada

4. Department of Biochemistry, Microbiology, and Immunology
   Faculty of Medicine
   University of Ottawa
   Ottawa, ON, Canada

5. Corresponding author:
   Sprott Center for Stem Cell Research
   Regenerative Medicine Program
   Ottawa Hospital Research Institute
   501 Smyth Road
   Ottawa, ON
   Canada K1H 8L6
  

**This PDF file includes:**
Supplemental legends and Figs. S1 to S6 and Tables S1 and S2.

**SUPPLEMENTAL FIGURES**

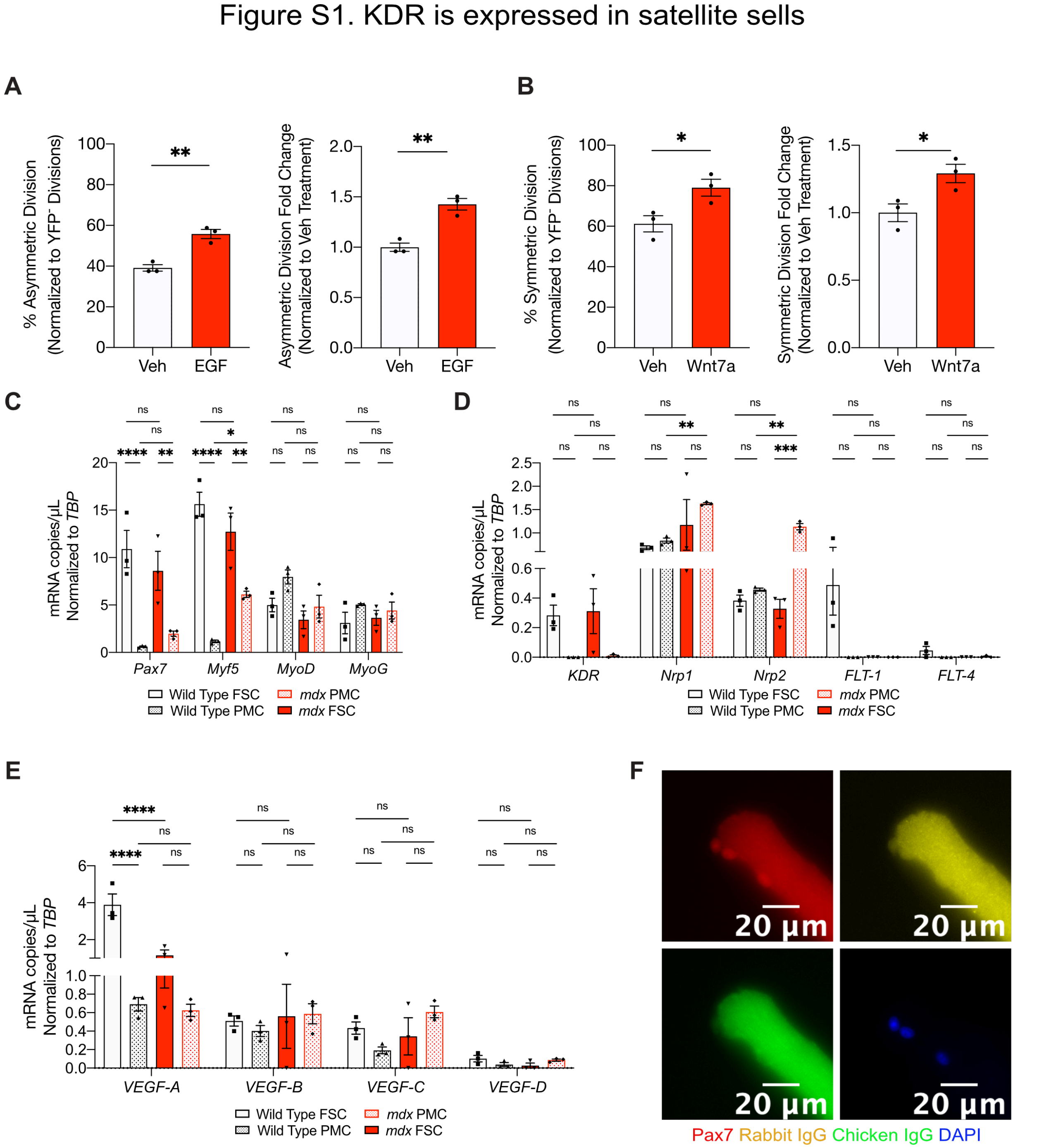

**Figure S1. KDR is expressed in satellite cells, Related to Figure 1**

(A) Validation of positive control EGF ligand on satellite stem cell asymmetric division on FDB myofibers normalized to total number of eYFP$^-$-containing divisions. Data is presented as mean. Error bars are ± SEM (n=3). ** p<0.01.

(B) Validation of positive control Wnt7a ligand on satellite stem cell symmetric division on FDB myofibers normalized to total number of eYFP$^-$-containing divisions. Data is presented as mean. Error bars are ± SEM (n=3). * p<0.05.

(C) Expression of myogenic regulatory factors (MRFs) normalized to *TBP*. Data is presented as mean. Error bars are ± SEM (n=3). ***p<0.001, **** p<0.0001.

(D) Receptor expression of *KDR* (VEGFR2), *Nrp1*, *Nrp2*, *FLT-1* (*VEGFR1*), *FLT-4* (VEGFR3) normalized to *TBP*. Data is presented as mean. Error bars are ± SEM (n=3). **** p<0.0001.

(E) Ligand expression of *VEGFA*, *VEGFB*, *VEGFC*, *VEGFD* normalized to *TBP*. Data is presented as mean. Error bars are ± SEM (n=3). **** p<0.0001.

(F) Pax7$^+$ (red) satellite cells stained with IgG controls for KDR (yellow), GFP (green) and nuclear DAPI (blue).

Figure S2. Validation of siRNA-mediated KDR knockdown

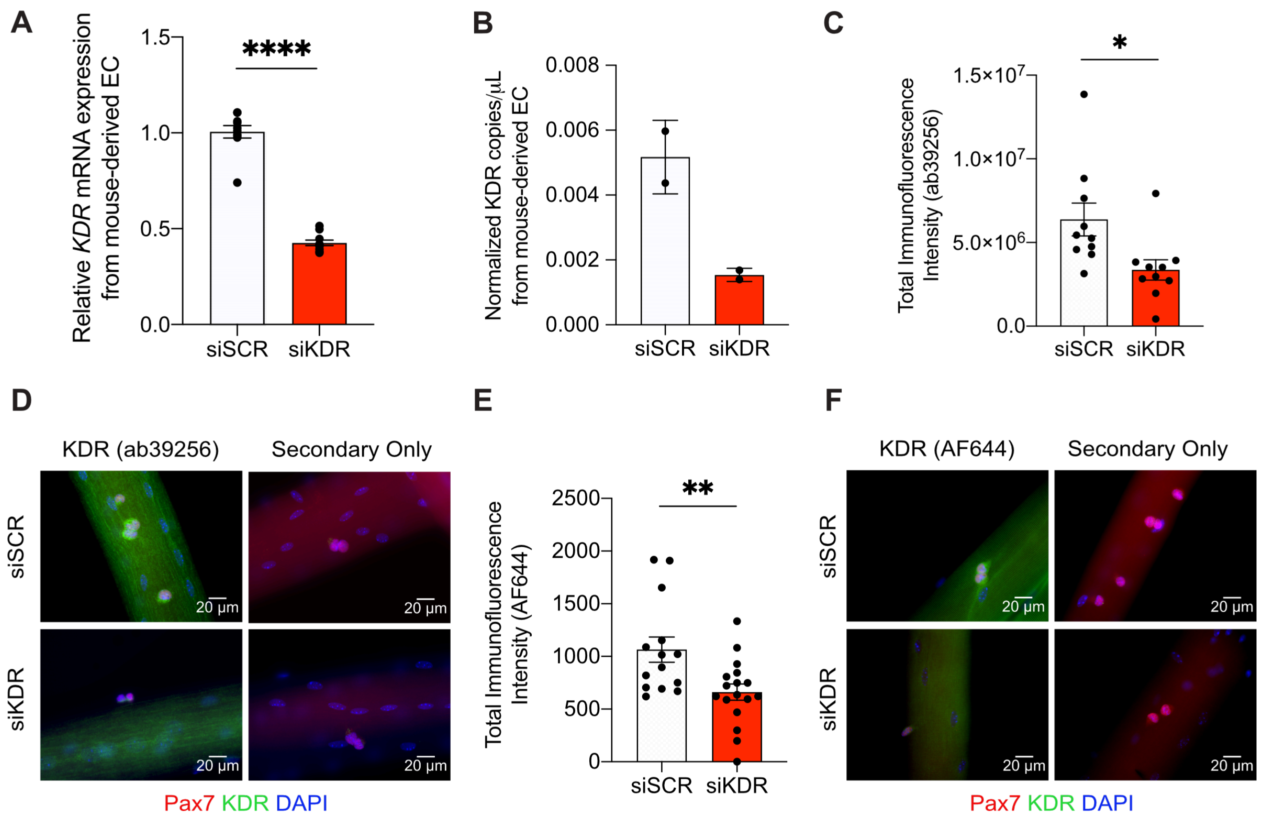

**Figure S2. KDR modulates satellite stem cell asymmetric division, Related to Figure 2**

(A) Validation of siKDR (10nM) on wild type mouse-derived endothelial cells (EC) by qPCR. Data is presented as mean. Error bars are ± SEM (n=5 technical replicates). **** $p<0.0001$.

(B) Validation of siKDR (10nM) on wild type mouse-derived endothelial cells (EC) by ddPCR using pre-validated KDR probes. Data is presented as mean. Error bars are ± SD (n=2).

(C) Validation of KDR antibody (ab39256) by siKDR (10nM) knockdown. Data is presented as mean. Error bars are ± SEM (n=10 cells per condition). *$p<0.05$

(D) Representative immunofluorescence (IF) images of Pax7$^{+}$ (red) satellite cells, KDR (green, ab39256), secondary only controls (Alexa Fluor 488) and DAPI (blue) nuclear stain on EDL single myofibers isolated from wild-type mice treated with siSCR control or siKDR.

(E) Validation of KDR antibody (AF644) by siKDR (10nM) knockdown. Data is presented as mean. Error bars are ± SEM (n=14-17 cells per condition) **$p<0.01$.

(F) Representative immunofluorescence (IF) images of Pax7$^{+}$ (red) satellite cells, KDR (green, AF644), secondary only controls (Alexa Fluor 488) and DAPI (blue) nuclear stain on EDL single myofibers isolated from wild-type mice treated with siSCR or siKDR.

Figure S3. KDR signaling requires the dystroglycan complex

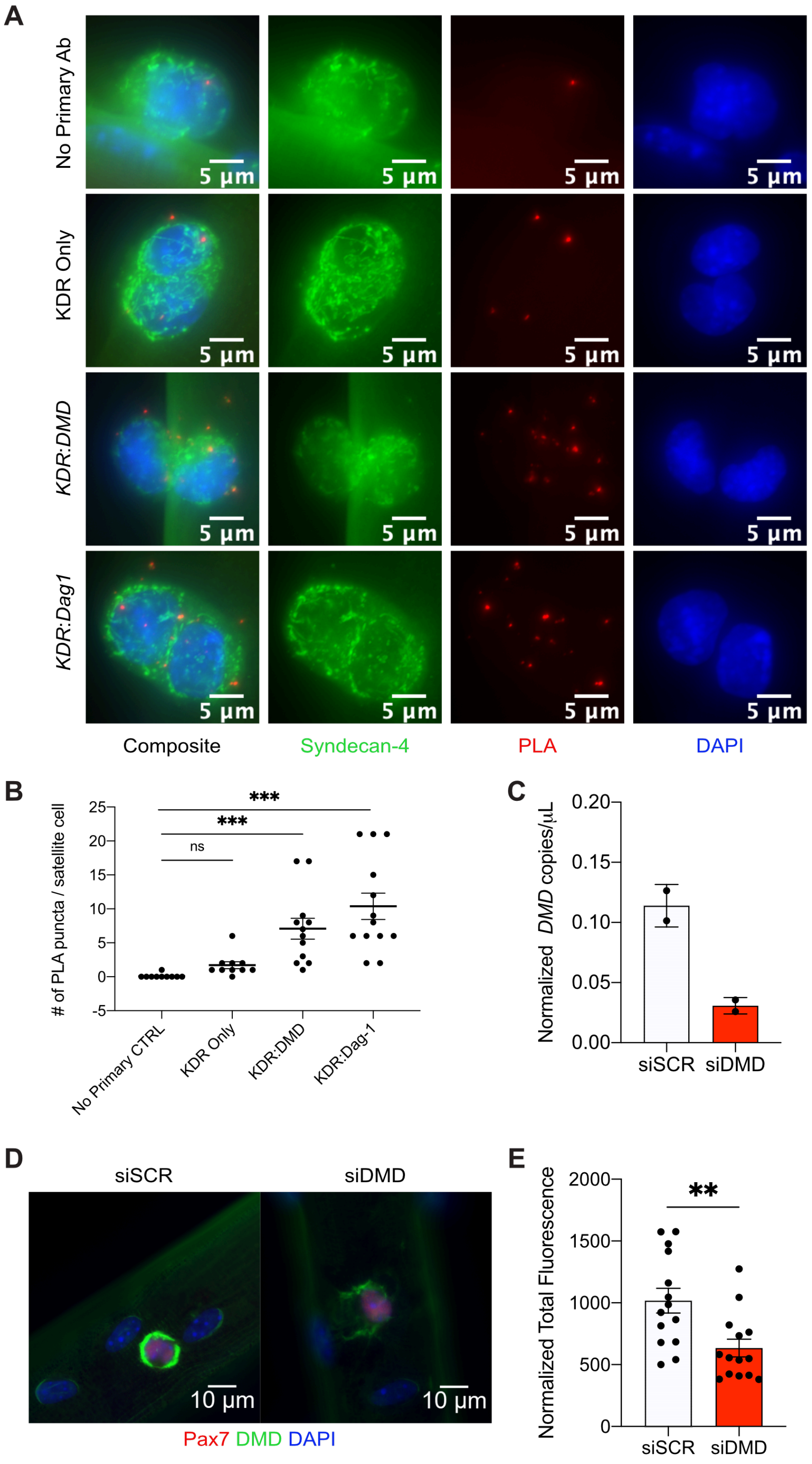

**Figure S3. MuSC KDR signaling requires the dystroglycan complex, Related to Figure 3**

(A) Representative immunofluorescence (IF) images of proximity ligand assay (PLA) punctate signal (red) of *KDR:Dag1* and *KDR:DMD* antibodies on wild-type Syndecan-4$^{+}$ (green) satellite cells and DAPI (blue) nuclear stain. Control conditions are secondary PLA probes only (No Primary Ab) or single primary KDR antibody (KDR only).

(B) Quantification of PLA signal in single primary antibody controls or *KDR:DMD* and *KDR:Dag1*. Data is presented as mean. Error bars are ± SEM (n=11-13 cells per condition). **p<0.001, ***p<0.0001.

(C) Validation of siDMD on wild-type myoblasts in 2-Day differentiation medium treated with either control (siSCR) or siDMD (10nM) by ddPCR using pre-validated DMD probes. Data is presented as mean. Error bars are ± SD (n=2).

(D) Representative immunofluorescence (IF) images of $Pax7^+$ (red) satellite cells, DMD (green) and DAPI (blue) nuclear stain on EDL single myofibers isolated from wild-type mice treated with siSCR (10μM) control or siDMD (10nM).

(E) Quantification of D and validation of siDMD (10nM) by immunofluorescence. Data is presented as mean. Error bars are ± SEM (n=14 cells per condition). **p<0.001.

### Figure S4. MuSC KDR-KO asymmetric division deficits limit satellite cell progenitors *in vivo*

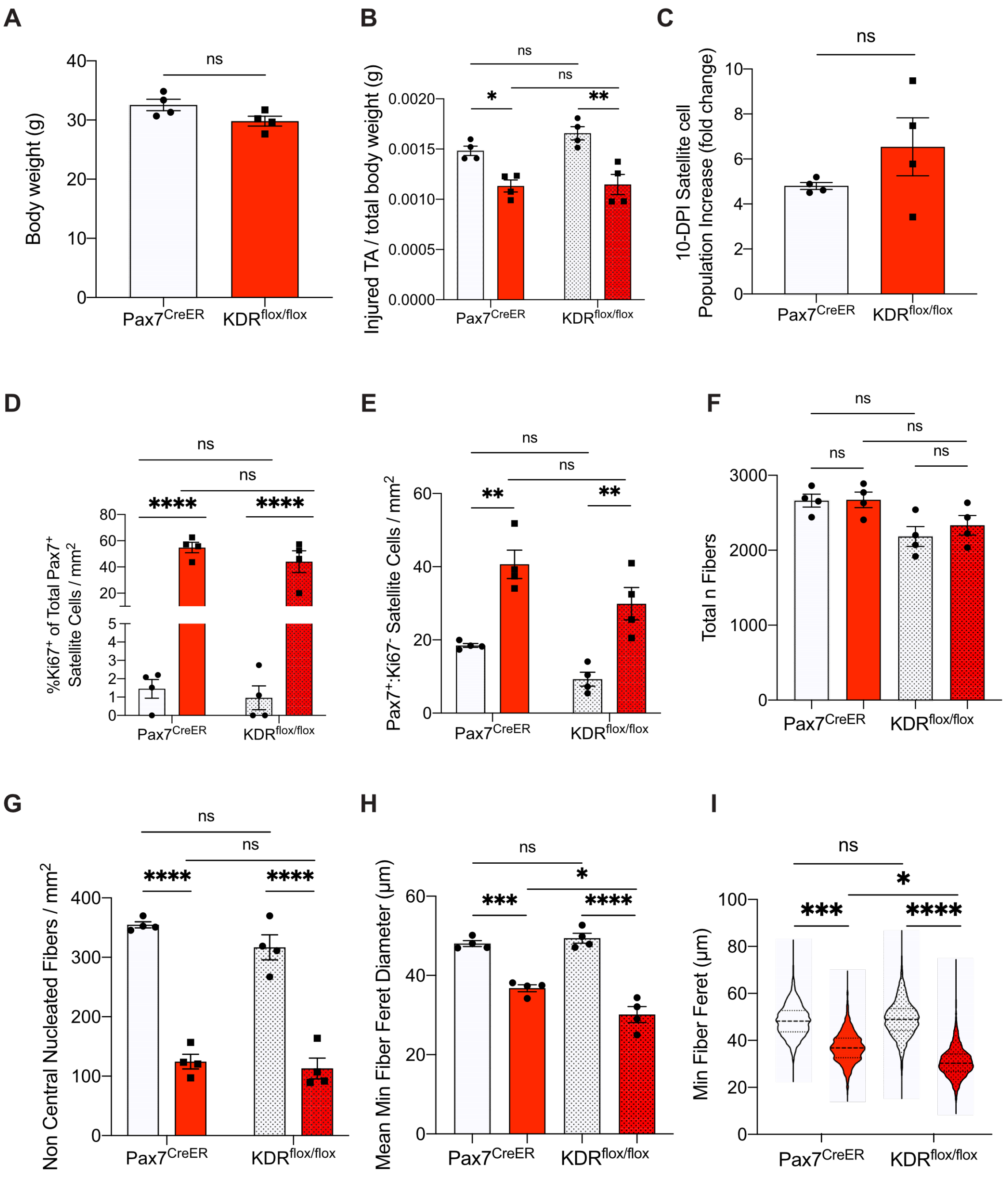

**Figure S4. MuSC KDR-KO asymmetric division deficits limits satellite cell progenitors *in vivo*, Related to Figure 5**

(A) Body weights (g) of wild type *Pax7$^{CreER}$:KDR$^{wt/wt}$* and wild type *Pax7$^{CreER}$:KDR$^{flox/flox}$* mice (n=4).
(B) Ratio of Injured TA weight over total body weight (g) of wild type *Pax7$^{CreER}$:KDR$^{wt/wt}$* and wild type *Pax7$^{CreER}$:KDR$^{flox/flox}$* mice (n=4).

(C) Fold change of satellite cell numbers in 10-days post injured TA compared to uninjured TA of wild type *Pax7$^{CreER}$:KDR$^{wt/wt}$* and wild type *Pax7$^{CreER}$:KDR$^{flox/flox}$* mice (n=4).

(D) Proportion of Pax7$^{+}$Ki67$^{+}$ satellite cells normalized to area in uninjured vs injured TA from wild type *Pax7$^{CreER}$:KDR$^{wt/wt}$* and wild type *Pax7$^{CreER}$:KDR$^{flox/flox}$* mice. Data is presented as mean. Error bars are ± SEM (n=4). **** p<0.0001.

(E) Quantification of Pax7$^{+}$Ki67$^{-}$ satellite cells normalized to area in uninjured vs injured TA from wild type *Pax7$^{CreER}$:KDR$^{wt/wt}$* and wild type *Pax7$^{CreER}$:KDR$^{flox/flox}$* mice. Data is presented as mean. Error bars are ± SEM (n=4). **p<0.01.

(F) Total fiber count of TA sections in uninjured vs injured TA from wild type *Pax7$^{CreER}$:KDR$^{wt/wt}$* and wild type *Pax7$^{CreER}$:KDR$^{flox/flox}$* mice. Data is presented as mean. Error bars are ± SEM (n=4).

(G) Quantification of non central nucleated fibers normalized to area in uninjured vs injured TA from wild type *Pax7$^{CreER}$:KDR$^{wt/wt}$* and wild type *Pax7$^{CreER}$:KDR$^{flox/flox}$* mice. Data is presented as mean. Error bars are ± SEM (n=4). ****p<0.0001.

(H) Mean minimum fiber ferret diameter (µm) of each biological replicate. Data is presented as mean. Error bars are ± SEM (n=4). *p<0.05, ***p<0.001, ****p<0.0001.

(I) Distribution of minimum fiber ferret (µm) in uninjured vs injured TA from wild type *Pax7$^{CreER}$:KDR$^{wt/wt}$* and wild type *Pax7$^{CreER}$:KDR$^{flox/flox}$* mice. Error bars are ± SEM (n=4). *p<0.05, ***p<0.001, ****p<0.0001.

Figure S5. MuSC KDR-KO result in reduced satellite cell numbers

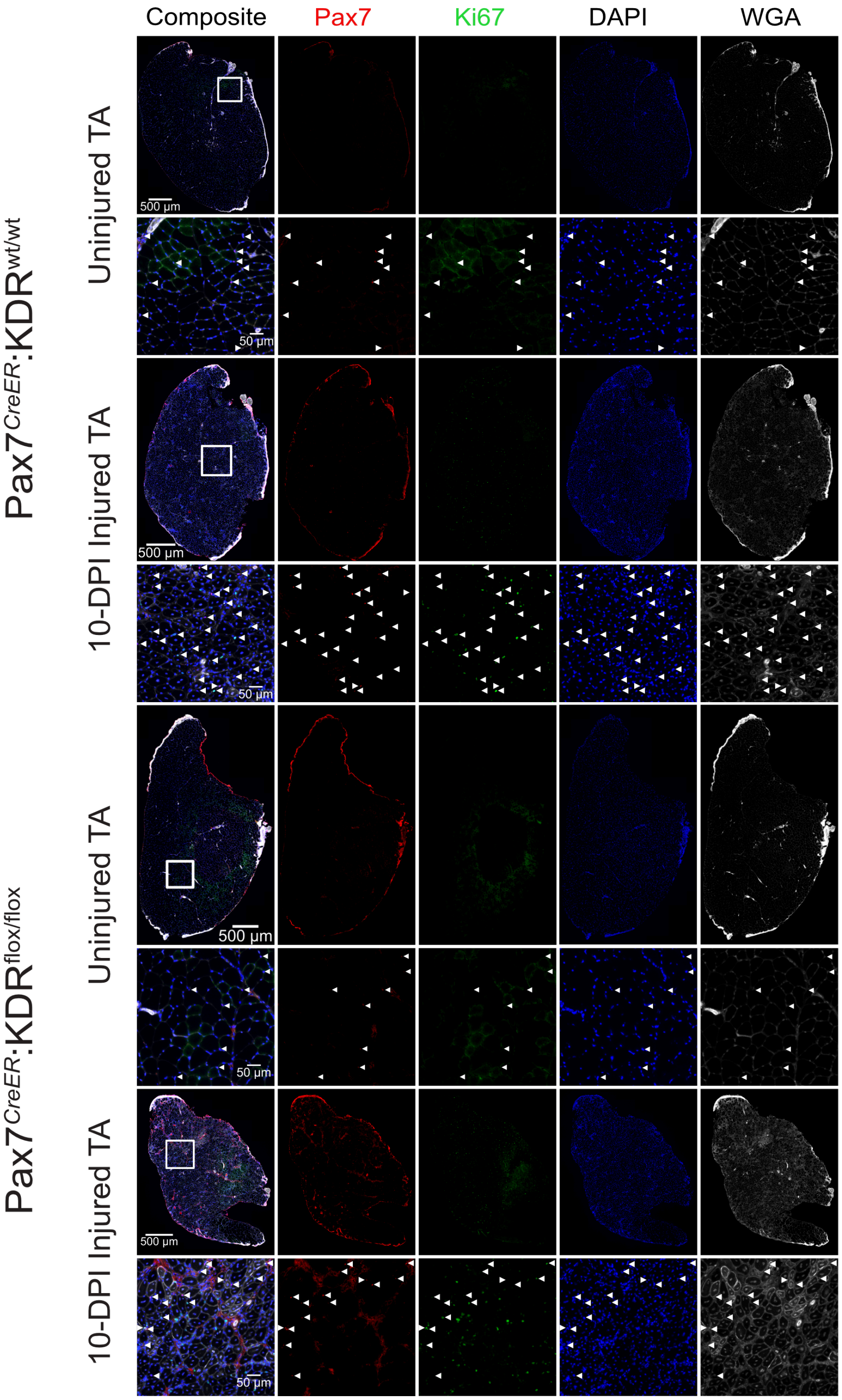

**Figure S5. MuSC KDR-KO result in reduced satellite cell numbers, Related to Figure 6**

Representative immunofluorescence (IF) images of uninjured and 10-DPI injured TA muscle sections from wild type *Pax7$^{CreER}$:KDR$^{wt/wt}$* and wild type *Pax7$^{CreER}$:KDR$^{flox/flox}$* mice showing Pax7$^{+}$ (red), Ki67$^{+}$ (green), Wheat Germ (white), DAPI (blue). Arrows indicate Pax7$^{+}$ cells.

Figure S6. MuSC KDR-KO perturbs muscle regeneration

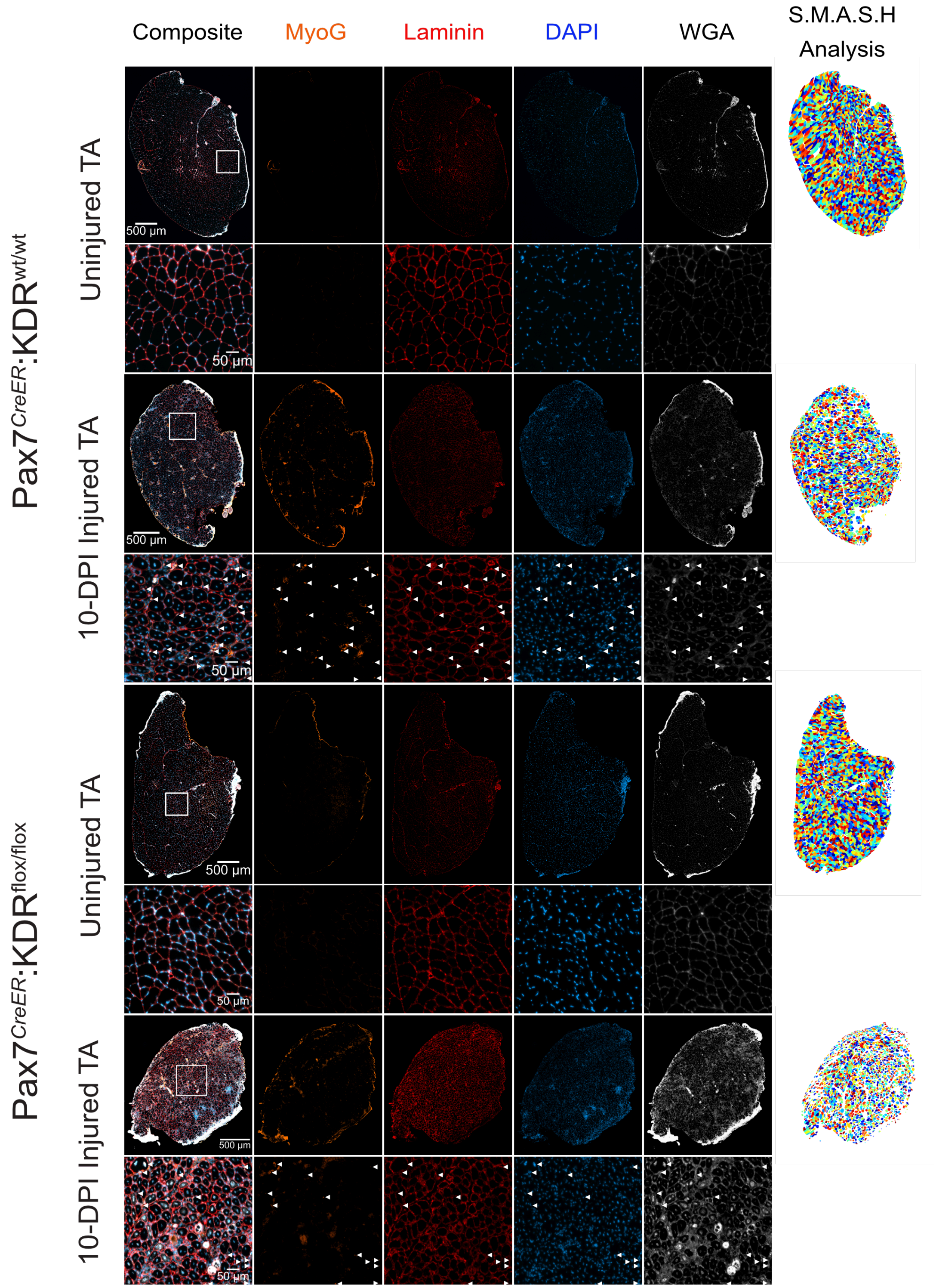

**Figure S6. MuSC KDR-KO perturbs muscle regeneration, Related to Figure 6**

Representative immunofluorescence (IF) images of uninjured and 10-DPI injured TA muscle sections from wild type $Pax7^{CreER}$:$KDR^{wt/wt}$ and wild type $Pax7^{CreER}$:$KDR^{flox/flox}$ mice showing $MyoG^{+}$ (orange), Laminin (red), Wheat Germ (white), DAPI (blue). Arrows indicate $MyoG^{+}$ cells. Far right column contains representative SMASH (semi-automatic muscle analysis)-generated mask of TA cross-sections used for analysis of fiber ferret diameter.

### SUPPLEMENTAL TABLES

**Table S1. Related to STAR Methods. siRNA Targeting Sequences**

| **siRNA Targeting Sequences (Figures 2, 3)** | |
|---|---|
| *KDR* (*VEGFR2*)<br>Integrated Life Technologies (Cat# mm.Ri.Kdr.13.1) | CUGCAAGUUUGGAAACCUAUCAACT<br>AGUUGAUAGGUUUCCAAACUUGCAGAA |
| *KDR* (*VEGFR2*)<br>Integrated Life Technologies (Cat# mm.Ri.Kdr.13.2) | AAUCAGUCAUUAUCUCCAGAACAGT<br>ACUGUUCUGGAGAUAAUGACUGAUUCC |
| *KDR* (*VEGFR2*)<br>Integrated Life Technologies (Cat# mm.Ri.Kdr.13.3) | GCAAAAGACUUUCAAACAGUGGCTC<br>GAGCCACUGUUUGAAAGUCUUUUGCUU |
| *DMD*<br>Integrated Life Technologies (Cat# mm.Ri.Dmd.13.1) | AAGAAGCUAGAACAAUCAUUACUGA<br>UCAGUAAUGAUUGUUCUAGCUUCUUGA |
| *DMD*<br>Integrated Life Technologies (Cat# mm.Ri.Dmd.13.2) | GUAUCAAACCAACAUCAUUACCUTT<br>UCAUCUGAUGUUCUAUUGCAGAAACAU |
| *DMD*<br>Integrated Life Technologies (Cat# mm.Ri.Dmd.13.3) | AAAGGUAAUGAUGUUGGUUUGAUAUC<br>GUUUCUGCAAUAGAACAUCAGAUGA |

**Table S2 Related to STAR Methods. Primers used for RT-qPCR and ddPCR**

| | | |
|---|---|---|
| **For RT-qPCR (Figure S2)** | | |
| *KDR* | Qiagen CAT#QT00097020 | Proprietary sequence |
| *PPIA* | FWD | CACTGCCAAGACTGAATG |
| *PPIA* | Rev | GTCGGAAATGGTGATCTTC |
| *RSP18* | FWD | AACGGTCTAGACAACAAGCTG |
| *RSP18* | Rev | AGTGGTCTTGGTGTGCTGAC |
| *GAPDH* | FWD | CCCAGA AGACTGTGGATGG |
| *GAPDH* | Rev | ACACATTGGGGGTAGGAACA |
| **For ddPCR (Figures 1 and S1)** | | |
| *Pax7* | Primer 1 CAT#Mm.PT.58.9286978 | TCCCCAGGATGATGAGACC |
| *Pax7* | Primer 2 CAT#Mm.PT.58.9286978 | TGTGACGGATGTGGTTCG |
| *Pax7* | Probe CAT#Mm.PT.58.9286978 | 56-FAM/TTGATGAAG/ZEN/ACCCCACCAAGCTGAT/3IABkFQ |
| *Myf5* | Primer 1 CAT#Mm.PT.58.5271235 | CACCTCCAACTGCTCTGAC |
| *Myf5* | Primer 2 CAT#Mm.PT.58.5271235 | ACATGCATTTGATACATCAGGAC |
| *Myf5* | Probe CAT#Mm.PT.58.5271235 | 5HEX/TGCCTGAA/ZEN/GTAACAGCCCTGTCTG/3IABkFQ |
| *MyoD* | Primer 1 CAT#Mm.PT.58.8193525 | GCTCTGATGGCATGATGGAT |
| *MyoD* | Primer 2 CAT#Mm.PT.58.8193525 | GACACAGCCGCACTCTT |
| *MyoD* | Probe CAT#Mm.PT.58.8193525 | 56-FAM/ACGACACCG/ZEN/CCTACTACAGTGAGG/3IABkFQ |
| *MyoG* | Primer 1 CAT#Mm.PT.58.6732917 | CTTGCTCAGCTCCCTCAAC |

| | | |
|---|---|---|
| *MyoG* | Primer 2 CAT#Mm.PT.58.6732917 | GACCGAACTCCAGTGCATT |
| *MyoG* | Probe CAT#Mm.PT.58.6732917 | 5HEX/AGCCCATGG/ZEN/THCCCAHTHAAT/3IABkFQ |
| *DMD* | Primer 1 CAT#Mm.PT.58.9899092 | GCTTATGTTGCCACCTCTGA |
| *DMD* | Primer 2 CAT#Mm.PT.58.9899092 | CTTCCGTCTCCATCAATGAACT |
| *DMD* | Probe CAT#Mm.PT.58.9899092 | 56-FAM/CAGAGCCCC/ZEN/TATCCTTCACAGCAT/3IABkFQ |
| *VEGFR1 (FLT-1)* | Primer 1 CAT#Mm.PT.58.43852013 | CAGAGCCAGGAACATATACACA |
| *VEGFR1 (FLT-1)* | Primer 2 CAT#Mm.PT.58.43852013 | AGGTCGTAGAGCCACTGAT |
| *VEGFR1 (FLT-1)* | Probe CAT#Mm.PT.58.43852013 | 5HEX/CTCGTTAGA/ZEN/GATTCGGAAGCGCCA/3IABkFQ |
| *KDR (VEGFR2)* | Primer 1 CAT#Mm.PT.58.5869721 | GGAATTGACAAGACAGCGACT |
| *KDR (VEGFR2)* | Primer 2 CAT#Mm.PT.58.5869721 | GGATCTTGAGTTCAGACATGAGG |
| *KDR (VEGFR2)* | Probe CAT#Mm.PT.58.5869721 | 56-FAM/AAGAAGGAG/ZEN/CAACACACAGCGAGC/3IABkFQ |
| *VEGFR3 (FLT-4)* | Primer 1 CAT#Mm.PT.58.13229320 | CTGAAGGATGGCACTCGAAT |
| *VEGFR3 (FLT-4)* | Primer 2 CAT#Mm.PT.58.13229320 | AGGTCAGAGAAAGCAGGTCT |
| *VEGFR3 (FLT-4)* | Probe CAT#Mm.PT.58.13229320 | 56-FAM/TGATGTGGC/ZEN/GTATGGCAGGAGTG/3IABkFQ |
| *Nrp1* | Primer 1 CAT#Mm.PT.58.9810806 | CCACAGAGAAGCCAACCATT |

| | | |
|---|---|---|
| *Nrp1* | Primer 2<br>CAT#Mm.PT.58.9810806 | GCAGAATGTCTTGTGAGAGC |
| *Nrp1* | Probe<br>CAT#Mm.PT.58.9810806 | 5HEX/AGCACCATC/ZEN/CAATCAGAGTTC<br>CCG/3IABkFQ |
| *Nrp2* | Primer 1<br>CAT#Mm.PT.58.30988321 | ATTATCCTGCCCAGCTATGAC |
| *Nrp2* | Primer 2<br>CAT#Mm.PT.58.30988321 | AGTGCTTATCCGAATGTCATCG |
| *Nrp2* | Probe<br>CAT#Mm.PT.58.30988321 | 5HEX/CCCTTCCCT/ZEN/ATCACTCCCTCG<br>AACA/3IABkFQ |
| *VEGFA* | Primer 1<br>CAT#Mm.PT.58.14200306 | CCGAAACCATGAACTTTCTGC |
| *VEGFA* | Primer 2<br>CAT#Mm.PT.58.14200306 | GACTTCTGCTCTCCTTCTGTC |
| *VEGFA* | Probe<br>CAT#Mm.PT.58.14200306 | 56-<br>FAM/TGCTGTACC/ZEN/TCCACCATGCCAA<br>G/3IABkFQ |
| *VEGFB* | Primer 1<br>CAT#Mm.PT.58.16515314 | CAAGTCCGAATGCAGATCCTC |
| *VEGFB* | Primer 2<br>CAT#Mm.PT.58.16515314 | GCTTCACAGCACTCTCCTT |
| *VEGFB* | Probe<br>CAT#Mm.PT.58.16515314 | 56-<br>FAM/TCCCTGGAA/ZEN/GAACACAGCCAAT<br>GT/3IABkFQ |
| *VEGFC* | Primer 1<br>CAT#Mm.PT.58.10862583 | GAAGTTCCACCATCAAACATGC |
| *VEGFC* | Primer 2<br>CAT#Mm.PT.58.10862583 | CAGCGGCATACTTCTTCACTA |
| *VEGFC* | Probe<br>CAT#Mm.PT.58.10862583 | 56-<br>FAM/CAATGCTTC/ZEN/AGTCGATTCGCAC<br>AGG/3IABkFQ |
| *VEGFD* | Primer 1<br>CAT#Mm.PT.58.43645237 | CACCTCCTACATCTCCAAACA |

| | | |
|---|---|---|
| *VEGFD* | Primer 2<br>CAT#Mm.PT.58.43645237 | CAAGCACTTACAACCCGTATG |
| *VEGFD* | Probe<br>CAT#Mm.PT.58.43645237 | 56-<br>FAM/TCAGTGCCT/ZEN/CTGACATCAGTGC<br>C/3IABkFQ |
| *TBP* | Primer 1<br>CAT#Mm.PT.58.17504874 | CGTGAATCTTGGCTGTAAACTTG |
| *TBP* | Primer 2<br>CAT#Mm.PT.58.17504874 | GTCCGTGGCTCTCTTATTCTC |
| *TBP* | Probe<br>CAT#Mm.PT.58.17504874 | 5HEX/ATCCCAAGC/ZEN/GATTTGCTGCAG<br>TC/3IABkFQ |

### KEY RESOURCES TABLE

| REAGENT or RESOURCE | SOURCE | IDENTIFIER |
|---|---|---|
| **Antibodies** | | |
| Mouse monoclonal anti-Pax7 | DSHB, Karakami et al., 1991 | RRID:AB_528428; https://doi.org/10.1016/S0925-4773(97)00097-X |
| Chicken polyclonal anti-GFP | Abcam | CAT#Ab13970; RRID:AB_300798 |
| Goat polyclonal anti-KDR | R&D Systems | CAT#AF644; RRID:AB_355500 |
| Rabbit polyclonal anti-KDR | Abcam | CAT#ab39256; RRID:AB_88343 |
| Rabbit polyclonal anti-DMD | Abcam | CAT#ab15277; RRID:AB_301813 |
| Rabbit polyclonal anti-Dag1 | Sigma | CAT#D1945; RRID:AB_1847470 |
| Chicken Syndecan-4 | Cornelison et al., 2004 | https://doi.org/10.1101/gad.1214204 |
| Rabbit polyclonal anti-Ki67 | Abcam | CAT#ab15580; RRID:AB_443209 |
| Mouse monoclonal anti-MyoG | R&D Systems | CAT#MAB66861; RRID:AB_10973343 |
| Rat monoclonal anti-Laminin-2 (α-2 Chain) | Millipore Sigma | CAT#L0663; RRID:AB_477153 |
| 647 conjugated anti-mouse α7-integrin | AbLab | CAT#67-0010-05 |
| PE/Cy7 VCAM CD106 | Cedarlane | CAT#105719; RRID:AB_2214047 |
| BV421 Integrin β1 | BD Bioscience | CAT#564131; RRID:AB_2738613 |
| PE conjugated anti-mouse CD45 | eBioscience | CAT#12-0451-82; RRID:AB_465668 |
| PE conjugated anti-mouse CD11b | eBioscience | CAT#12-0112-81; RRID:AB_465546 |
| PE conjugated anti-mouse Sca-1 (Ly-6A/E) | BD Pharmingen | CAT#553108; RRID:AB_394629 |
| PE conjugated anti-mouse CD31 (PECAM1) | BD Pharmingen | CAT#553373; RRID:AB_394819 |
| Satellite Cell Isolation Kit | Miltenyi Biotec | CAT#130-104-268 |
| Anti-Integrin α7 | Miltenyi Biotec | CAT#130-104-261 |
| CD31 MicroBeads | Miltenyi Biotec | CAT#130-097-418 |
| **Biological Samples** | | |
| Mouse: WT *Myf5-Cre::R26R-eYFP* Single FDB myofibers | This paper | N/A |
| Mouse: WT *Myf5-Cre::R26R-eYFP* Single EDL myofibers | This paper | N/A |
| Mouse: *WT Pax7-nGFP* FACs sorted satellite cells | This paper | N/A |
| Mouse: *MDX Pax7-nGFP* FACs sorted satellite cells | This paper | N/A |
| Mouse: *WT* $Pax^{CreER}$::$KDR^{wt/wt}$ TA muscle | This paper | N/A |
| Mouse: *WT* $Pax^{CreER}$::$KDR^{flox/flox}$ TA muscle | This paper | N/A |
| **Chemicals, Peptides, and Recombinant Proteins** | | |

| | | |
|---|---|---|
| Ontario Institute for Cancer Research (OICR) Kinase Inhibitor Library | OICR | https://oicr.on.ca |
| 4',6-Diamidino-2-phenylindole dihydrochloride (DAPI) | Sigma-Aldrich | CAT#D8417 |
| Wheat Germ Agglutinin (WGA) Alexa Fluor 647 Conjugate | Life Technologies | CAT#W32466 |
| EGF Ligand | Miltenyi Biotec | CAT#130-093-825 |
| Wnt7a Ligand | R&D Systems | CAT#3008WN025/CF |
| VEGF E (Orf Virus) Ligand | ProSpec | CAT#CYT-263 |
| Click-iT EdU Alexa Fluor 647 Imaging Kit | Invitrogen | CAT#C10340 |
| Cardiotoxin Gamma (Latoxan) | Cedarlane | CAT# L8102 |
| Tamoxifen >99% | Sigma-Aldrich | CAT#T5648 |
| Teklad Custom Tamoxifen Diet | Envigo | CAT# TD.130857 |
| DMEM | Gibco | CAT#11320082 |
| Ham-F10 | Wisent | CAT#318-051-CL |
| Collagenase Type I | Sigma | CAT#C0130 |
| Fetal Bovine Serum | Wisent | CAT#098-150 |
| Chicken Embryo Extract | MP Biomedicals | CAT#92850145 |
| bFGF (human recombinant) | Cedarlane | CAT#GF003AFMG(CH) |
| Penicillin/Streptomycin (100x) | GIBCO | CAT#15140122 |
| Lipofectamine RNAimax | Life Technologies | CAT#11668019 |
| Critical Commercial Assays | | |
| Macherey-NagelSupplier Diversity Partner NUCLEOSPIN RNA II | Fisher Scientific | CAT#740955.250 |
| iScript™ cDNA Synthesis Kit | Bio-Rad | CAT#170-8891 |
| SsoFast ™ EvaGreen Supermix | Bio-Rad | CAT#172-5202 |
| PicoPure RNA Isolation | Life Technologies | CAT#KIT0204 |
| Duolink Detection Reagents Red | Sigma-Aldrich | CAT#DUO92008-100RXN |
| Duolink In Situ PLA Probe Anti-Goat PLUS | Sigma (Olink Bioscience | CAT#DUO92003-100RXN |
| Duolink In Situ PLA Probe Anti-Goat MINUS | Sigma (Olink Bioscience | CAT#DUO92006-100RXN |
| Duolink In Situ PLA Probe Anti-Rabbit PLUS | Sigma (Olink Bioscience | CAT#DUO92002-100RXN |
| Duolink In Situ PLA Probe Anti-Rabbit MINUS | Sigma (Olink Bioscience | CAT#DUO92005-100RXN |
| Neural Dissociation Kit P | Miltenyi Biotec | CAT#130-092-628 |
| Endothelial Cell Growth Medium MV2 Kit | Promocell | CAT#10175-25 |
| Deposited Data | | |
| Experimental Models: Cell Lines | | |
| Mouse: Passage <10 wild-type primary satellite cells | This paper | N/A |
| Mouse: Passage 5 wild-type primary endothelial cells | This paper | N/A |
| Experimental Models: Organisms/Strains | | |
| Mouse: Wild-type: C57BL/10ScSn | The Jackson Laboratory | JAX: 000476; RRID:IMSR_JAX:000476 |

| | | |
|---|---|---|
| Mouse: *MDX*: *DMD*$^{mdx}$-C57BL/10ScSn | The Jackson Laboratory | JAX: 001801; RRID:IMSR_JAX:001801 |
| Mouse: Pax7$^{CreER}$: *Pax7*$^{tm1(cre/ERT2)Gaka}$-B6.Cg | The Jackson Laboratory | JAX: 017763; RRID:IMSR_JAX:017763 |
| Mouse: Myf5$^{Cre}$: *Myf5*$^{tm3(cre)Sor}$-B6.129S4 | The Jackson Laboratory | JAX: 007893; RRID:IMSR_JAX:007893 |
| Mouse: R26R-*eYFP*: Gt*(ROSA)26Sor*$^{tm1(EYFP)Cos}$-B6.129X1 | The Jackson Laboratory | JAX: 006148; RRID:IMSR_JAX:006148 |
| Mouse: KDR$^{flox/flox}$: *KDR*$^{tm2Sato}$-B6.129S1 | The Jackson Laboratory | JAX: 018977; RRID:IMSR_JAX:018977 |
| Mouse: Pax7-*nGFP*: *Pax7*$^{tm1.2Tajb}$-B6.SJL | Sambasivan et al., 2009 | RRID:MGI:5495462; https://doi.org/10.1016/j.devcel.2009.05.008 |
| Oligonucleotides | | |
| DsiRNA KDR | Integrated DNA Technologies | CAT#mm.Ri.Kdr.13 |
| DsiRNA DMD | Integrated DNA Technologies | CAT#mm.Ri.Dmd.13 |
| Primers for siRNA, see Table S1 | Integrated DNA Technologies | N/A |
| KDR Quantitect Primer Assay | Qiagen | CAT#QT00097020 |
| KDR PrimeTime qPCR Probe Assay (ddPCR) | Integrated DNA Technologies | CAT#Mm.PT.58.5869721 |
| VEGFA PrimeTime qPCR Probe Assay (ddPCR) | Integrated DNA Technologies | CAT#Mm.PT.58.14200306 |
| DMD PrimeTime qPCR Probe Assay (ddPCR) | Integrated DNA Technologies | CAT#Mm.PT.58.9899092 |
| Primers for ddPCR, see Table S2 | Integrated DNA Technologies | N/A |
| Recombinant DNA | | |
| Software and Algorithms | | |
| Fiji - ImageJ | Schneider et al., 2012 | RRID:SCR_002285; https://imagej.nih.gov/ij/ |
| Zen Imaging Suite | ZEISS | RRID:SCR_013672; https://www.zeiss.com |
| Prism | GraphPad | RRID:SCR_002798; www.graphpad.com |
| HCS Studio Cell Analysis | Thermo Fisher Scientific | RRID:SCR_016787; https://www.thermofisher.com/ |
| MATLAB | Mathworks | RRID:SCR_001622; https://www.mathworks.com |

| | | |
|---|---|---|
| IMARIS | Oxford Instruments | RRID:SCR_007370; https://imaris.oxinst.com |
| SMASH | Smith and Barton, 2014 | https://doi.org/10.1186/2044-5040-4-21 |
| Other | | |
